# TIM3^+^ Tumor Associated M2 Macrophages Impair Antitumor T Cell Immunity and Promote Gastric Cancer Progression and Peritoneal Metastasis

**DOI:** 10.64898/2026.04.15.716939

**Authors:** Xiaodan Yao, Yibo Fan, Junsong Zhao, Yan-ting Yann Zhang, Dipti Athavale, Curt Balch, Mikel Ghelfi, Anthony Pompetti, Joseph Zhao, Ailing Scott, Jiankang Jin, Young Ki Hong, Jamin Morrison, Madeline Torres, Shilpa S Dhar, Linghua Wang, Jimmy Bok-Yan So, Patrick Tan, Raghav Sundar, Francis Spitz, Generosa Grana, Jaffer A Ajani, Shumei Song

**Author notes:** The authors declare no potential conflicts of interest. Corresponding author: Shumei Song, MD, Ph.D, Coriell Institute for Medical Research, New Jersey, 08103, USA.; Jaffer A. Ajani, MD, The University of Texas MD Anderson Cancer Center, Houston, TX 77030, USA.

## Abstract

Peritoneal metastases (PM) are the leading cause of cancer-related death in gastric cancer (GC) patients with survival typically < 9 months. Here, we demonstrate that TIM3 and its ligands are increased along the GC continuum and associated with poor survival. Integrated omics analyses and functional studies revealed highly enriched TIM3 in CD163^+^ tumor associated M2 immunosuppressive macrophages significantly promote tumor cell invasion and tumor growth *in vivo,* while TIM3 depletion in macrophages reduced tumor cell malignant attributes and increased T cell immunity from PBMCs or CD45+ immune cells of malignant ascites in co-culture system. By cytokine and kinase arrays, we discovered that depletion of TIM3 in macrophages reduced the production of notable secretome of cytokines/chemokines from M2 macrophages; and the protumor function of TIM3^+^ macrophages rely on the p90RSK1/2/CCL20 axis. Finally, we reveal that TIM3 blockage or genetic KO had superior antitumor activity in combination with anti-PD1 immunotherapy and mitomycin C (MMC) chemotherapy. Together, this study uncovers an important role for TIM3 in tumor associated M2 macrophages and underscores the potential of TIM3 blockage in GC patients with PM.

**Statement of significance:** In this study, we show TIM3 increases along GC continuum, and highly enriched on tumor associated M2 macrophages that fuel tumor growth; and suppress T cell function via p90RSK1/2/CCL20 axis. TIM3 depletion restores T-cell immunity and curbs tumor growth. TIM3 blockade combined with anti-PD1 and mitomycin C provide a novel therapeutic strategy for GC patients with PM.

## Introduction

Gastric cancer (GC) is a major global health burden. In the U.S, 31,510 new cases, and 10,740 GC deaths are expected in 2026^1^. In general, over 90% of all cancer-associated deaths are caused by metastases, and over 40% of GC patients develop peritoneal metastases (PM; malignant ascites or implants in peritoneal cavity), with dismal prognosis and short survival (less than 9 months).^2–6^ Such patients have a poor quality of life, and are difficult to manage^2, 7^, with no effective targeted or immune therapy available. Currently available treatment trials are hyperthermic intraperitoneal chemotherapy (HIPEC, e.g., mitomycin C and paclitaxel at 41°C – 42°C) for GC patients with PM (GCPM). However, this has not been efficacious in GCPM patients, underscoring an urgent need to better understand this disease at the molecular level for exploring novel immune therapies and tumor cell-based targeted therapies.

T-cell immunoglobulin and mucin domain 3 (TIM-3), a member of the T cell immunoglobin and mucin domain protein family, was initially identified on the surface of T helper 1 (Th1) cells and cytotoxic (CTL) lymphocytes^8^ as an immunosuppressive molecule through interacts with its ligands (GAL-9, CEACAM1, HMGB1, and PtdSer)^9^; and its expression correlates with nodal metastases and advanced cancer stage ^10, 11^. Expression of TIM3 on CD8 T cells in the tumor microenvironment (TME) is considered a cardinal sign of T cell dysfunction^12^. However, TIM3 is also expressed on several other types of immune cells including DCs, macrophages, FoxP3+ Tregs, etc.^13, 14^ ^15, 16^, and one recent study reported that TIM3 conditional deletion on DCs, but not on CD4 or CD8 cells, promoted strong anti-tumor immunity^15^. A more recent study identified that TIM3 was also expressed in glioma cells in addition to T cells and several other immune cells, supporting it as a bona fide target in diffuse intrinsic pontine glioma (DIPG)^17^. However, the functional role of TIM3 on macrophages in promoting GCPM and its potential therapeutic or clinical translation in advanced GC remain unknown.

In this study, using multiple reverse translational approaches, we identified that TIM3 is highly enriched in tumor associated M2 macrophages in GCPM. TIM3 knockout (KO) in macrophages impaired tumor cell malignant behaviors in a co-culture *in vitro* system and suppressed tumor growth induced by M2 macrophages *in vivo*; while depletion of TIM3 in macrophages significantly increased CD3 and CD8 T cell immunity. Further, cytokine arrays revealed that TIM3 dictates the secretome of M2 macrophages, TIM3 KO in macrophages eliminated the production of notable cytokines (i.e. CCL1, CCL4, CCL5, CCL7, IL-1b, IL-6, and CCL20) that promote metastases. Among these, CCL20 expression and secretion was significantly reduced upon TIM3 depletion; and both CCL20 and its receptor CCR6 are highly increased in tumors compared to normal tissues and associated with poor survival. Concordantly, recombinant CCL20 (rCCL20) can rescue effects of TIM3 KO in macrophages on tumor cells and T cell immunity indicating that oncogenic functions of macrophage associated TIM3 occur partially through the CCL20/CCR6 axis. Furthermore, kinase array and functional study in U937 macrophages with TIM3 KO vs control revealed that p90RSK1/2 is the key kinase mediated TIM3 regulation of CCL2 in M2 macrophages. Most importantly, we revealed that the anti-tumor effects of TIM3 blockage depend on macrophages, and abrogation of TIM3 enhances the efficacy of immune checkpoint blockade (anti-PD1), or MMC chemotherapy in PDXs and a syngeneic mouse model.

## METHOD DETAILS

### Cell lines and culture

The patient-derived GCPM cell line GA0518 and its derivative GA0518mCherry Luciferase were generated from malignant ascites of a GC patient with peritoneal metastasis, as described previously^22^. These cells were cultured in 7% fetal bovine serum (FBS, HyClone, cytiva)-RPMI medium. Murine KP-Luc2 GC cells were reported previously^20^ and cultured in 7% FBS-DMEM (Gibco). Human monocyte U937 and murine macrophage RAW264.7 were purchased from ATCC, and both monocytes and macrophages were cultured in 7% FBS-RPMI. All cell lines were authenticated and confirmed to be mycoplasma-free every six months.

### Generation of *TIM3* KO cells from human U937 and murine RAW264.7 macrophage using LentiCRISPR/CAS9

The CRISPRs (clustered regularly interspaced short palindromic repeats) /Cas9 system has been used to KO the *TIM3* gene in U937 and RAW267.4 (RAW) cells. Guide RNAs (gRNA) design follows the principle in MIT website http://crispr.mit.edu/. The gRNA sequences were cloned into the pLentiCRISPR v2 mOrange (V2mO) vector.

Complementary oligonucleotides for each gRNA were annealed by heating to 100°C to form duplexes, which were then ligated into BsmBI-digested V2mO using T4 DNA ligase (NEB, Ipswich, MA). Recombinant lentiviral plasmids containing the target gene were co-transfected with packaging plasmids pCMV-dr8.2 and pCMV-VSV-G at a ratio of 10:10:1 into HEK293T cells at approximately 70% confluency using JetPRIME transfection reagent (Polyplus, France). Viral supernatants were harvested and used immediately for transduction. U937 and RAW264.7 cells were seeded in 6-well plates at ∼70% confluency and transduced with lentiviral supernatants in the presence of 8 µg/mL polybrene.

Transduced cells were selected with puromycin for 3–6 days, followed by single-cell cloning. Successful TIM3 knockout clones were identified by Western blot analysis using a TIM3-specific antibody.

#### Immune cell function analysis by flow cytometry

Fresh mouse tumor tissues were digested with 2 mg/mL collagenase IV (Gibco, 17104-019) and 2 mg/mL dispase (Gibco, 17015-041) at 37 °C for 30 minutes, followed by mechanical dissociation using a gentleMACS Dissociator (Miltenyi Biotec). The resulting single-cell suspensions were washed with PBS and stained with Live/Dead Aqua dye (1:1000; Life Technologies) in PBS. Cells were then incubated with a surface antibody cocktail containing anti-mouse CD3–APC-Cy7 (BioLegend, Cat#100222) and CD8-Alexa Fluor 700 (BioLegend, Cat#100730) in FACS buffer (PBS containing 0.5% BSA) at 4 °C for 30 minutes. After washing once with FACS buffer, cells were fixed and permeabilized, washed twice, and subjected to intracellular staining with IFNγ–PE-Cy7 (BioLegend, Cat#505825) in permeabilization buffer for 30 minutes in the dark on ice. Data were acquired on a Gallios 561 flow cytometer (Beckman Coulter) and analyzed using FlowJo software (version 10).

For in vitro assays, human or mouse PBMCs were co-cultured with U937 or RAW264.7 cells with or without TIM3 knockout for 2 days. Suspended cells were collected, washed with FACS buffer, and processed directly for flow cytometry. Surface staining was performed with CD3-APC-Cy7 (BioLegend, Cat#300318) and CD8-Alexa Fluor 700 (BD Biosciences, Cat#557945), followed by intracellular staining with Granzyme B BV421 (BD Biosciences, Cat#563388), Perforin PE-CF594 (BD Biosciences, Cat#563763), and IFNγ-PE-Cy7 (BioLegend, Cat#506518). Flow cytometry data were acquired using a BD FAC Symphony A3. All experiments were performed in biological triplicate and repeated at least three times.

#### Macrophage repolarization and immune analysis after co-culturing with PBMCs

U937 or RAW264.7 monocytic cells were treated with 80 nM phorbol 12-myristate 13-acetate (PMA; Sigma, Cat#16561-29-8) for 48 hours to induce differentiation into M1-like macrophages prior to co-culture. Human PBMCs were stimulated with 100 IU/mL human IL-2 (R&D Systems, Cat#202-IL-010/CF) for 2 days. After washing with PBS, 5 × 10□U937- or RAW264.7-derived M1 macrophages, with or without TIM3 knockout, were repolarized toward an M2-like state by incubation with conditioned medium from GA0518 tumor cells. PBMCs were then added and co-cultured in 12-well plates for an additional 48 hours. Following co-culture, PBMCs were collected and analyzed by flow cytometry to assess cytokine and cytotoxic marker expression (IFNγ, Granzyme B, and Perforin) in CD3□ and CD8□ T cells. Adherent U937 or RAW264.7 macrophages were harvested for analysis of M2-associated gene expression, including *CCL2* and *IL10*, by RT-qPCR using human- or murine-specific primers.

#### Matrigel Invasion Assay

The invasion assay was performed as described previously^20^. Briefly, human GA0518 and murine KP-Luc2 tumor cells (2.5 ×10^4^) were placed in ThinCerts-TC inserts with 0.5% FBS-RPMI medium, and co-cultured with human U937 or mouse RAW264.7 MΦ (7.5 ×10^4^) with or without TIM3 KO, in 24-well plates, for 2 days. The inserts were then fixed with 2% PFA-PBS, stained with 0.5% crystal violet blue, washed, their inner sides wiped out, and the blue transferred tumor cells observed under a microscope and counted.

#### ELISA

To measure CCL20 secretion, culture supernatants were collected from equal numbers of U937- or RAW264.7-derived macrophages with or without TIM3 knockout. Secreted CCL20 levels were quantified using a human/mouse CCL20 ELISA kit according to the manufacturer’s instructions (R&D Systems, DY360-05).

#### Cytokine Array

U937 control and U937 TIM3 knockout cells (5 × 10□ per well in 6-well plates) were treated with 100 nM PMA for 2 days to induce macrophage differentiation, followed by incubation with GA0518-conditioned medium (cGM) for an additional 2 days to promote repolarization toward a tumor-associated M2-like phenotype. Cells were then washed with PBS and cultured in normal growth medium (GM) for a further 24 hours. Culture supernatants were collected and analyzed using the RayBio® C-Series Human Cytokine Antibody Array C5 kit (RayBiotech, AHH-CYT-5-8) according to the manufacturer’s protocol.

#### Phosphorylation Kinase Array

U937 TIM3 knockout and control cells were polarized to an M1-like state using PMA and subsequently repolarized toward an M2-like macrophage phenotype by incubation with GA0518 tumor cell–conditioned medium. Cells were then harvested, and total protein lysates were prepared and quantified. Equal amounts of protein were applied to the Proteome Profiler Human Phospho-Kinase Array Kit (R&D Systems) to assess the phosphorylation status of 22 protein kinases, according to the manufacturer’s instructions. Array signals were scanned and imaged, and kinase spot intensities were quantified and compared using iBright software.

#### qRT-PCR

Total RNA was extracted from cultured cells with TRIzol, converted to cDNA using LinaScript RT SuperMix Kit, E3010L (New England BioLabs), and subjected to q-PCR with PowerUp SYBR Green Master Mix (Applied Biosystems A25742). Primers used in this study for q-PCR were listed in Supplemental Table 2.

#### Western Blot

Western blot analyzes were performed as previously described^39^. Briefly, the quantified protein samples were run for electrophoresis using SurePAGE Gels in Tris-MOPS-SDS Buffer (GenScript) and dry-electorally blotted to PVDF membrane (Invitrogen, iBolt 3 Transfer Stack). The membranes were bound with individual primary antibodies TIM-3/HAVCR2 antibody (Sino Biological), mouse anti-human/mouse □-actin (R&D), and antibodies from Cell Signaling Technology: Phospho-p90RSK1/2 (Ser380), Phospho-AKT (Ser473), Phospho-p44/42 MAPK (ERK1/2), anti-rabbit or anti-mouse linked with HRP in Pierce Protein-Free-Blocking Buffer (Thermo Scientific). Finally, the membranes were visualized with iBright 1500 (Invitrogen).

#### Immune killing assay

GA0518–mCherry–luciferase target tumor cells (T) were seeded at 2 × 10□ cells per well in 96-well plates and allowed to adhere overnight. U937-derived M2 macrophages (1 × 10□ cells per well) were then added under three conditions: (1) no U937 cells, (2) U937 control cells, and (3) U937 TIM3 knockout cells. Human effector PBMCs (E) were subsequently added at effector-to-target (E:T) ratios of 1:1, 4:1, and 16:1 in RPMI-1640 supplemented with 10% FBS and co-cultured for 48 hours. Tumor cell viability and cytotoxicity were quantified by measuring luciferase activity. All conditions were performed in triplicate.

#### Immunofluorescence (IF)

Immunofluorescence was performed as previously described^40^. Antigen retrieval was performed using Antigen Unmarking Solution (BioGenex laboratories, Cat# HK0805K-GP). Antibodies specific to human TIM3, CD163, human Ki67 (Fisher Scientific, Cat#RM-9106-S1, 1:200), human YAP1(Abcam, Cat# ab205270, 1:200) and SOX9 (Abcam, Cat# ab76997, 1:200) followed by secondary antibodies. Slides were then mounted with DAPI-containing Vecta shield Mounting Medium (Vector Laboratories, Cat# H-1200-10), visualized under the Nicon T2 confocal laser scanning microscope.

#### CyTOF Mass Cytometry

CyTOF was performed as previously described^40^. In brief, 1.5 × 10□ live ascites cells per sample were washed and blocked with 5% FBS, then resuspended in 0.5% BSA in PBS and incubated with cell surface antibodies at room temperature for 1 hour. Cells were subsequently stained with cisplatin (25 µM; Sigma) to discriminate dead cells, followed by fixation in 1.6% formaldehyde in PBS and permeabilization in 90% methanol at 4 °C overnight. After washing, cells were incubated with 50 µL of intracellular antibody cocktail for 1 hour at room temperature, followed by staining with 62.5 nM iridium intercalator (Fluidigm) at 4 °C overnight. Samples were acquired on a CyTOF2 mass cytometer (Fluidigm). Antibodies used for CyTOF analysis are listed in Supplemental Table 3. CyTOF data were pre-gated to exclude dead cells and irrelevant populations, then concatenated using FlowJo software (version 10). Dimensionality reduction using t-distributed stochastic neighbor embedding (t-SNE) and clustering analysis using PhenoGraph were performed with the open-source R package CyTOFkit (https://github.com/JinmiaoChenLab/cytofkit

#### Single Cell RNA Sequencing (scRNA-seq) Analysis

scRNAseq analyses were performed as previously described^35, 41^. Briefly, droplet-based 3′ single-cell RNA sequencing (10x Genomics) was conducted on peritoneal ascites samples (n = 20) at the SMF Core Facility, MD Anderson Cancer Center. Sequencing reads were aligned and transcripts were assigned to individual cells based on cellular barcodes using Cell Ranger. Downstream analyses were performed using Seurat in R (version 4.0). Genes detected in at least three cells and cells expressing at least 200 genes were retained for analysis. The first 24 principal components were used for clustering (resolution = 0.5) and for visualization by UMAP, dot plots, and violin plots.

#### KP-Luc2 Syngeneic mouse models

All in vivo animal studies were approved by the Institutional Animal Care and Use Committee (IACUC; protocol ID: 23-016) at Cooper University Hospital. Both male and female mice were used in these studies, and sex was considered as a biological variable. All *in vivo* studies were conducted with mice randomly assigned to experimental groups.

##### TIM3^+^ tumor associated M2 Macrophage–Mediated Tumor Growth in vivo (see Figure 3D-3F)

To evaluate the role of TIM3^+^ M2 macrophages in tumor progression, mouse KP-Luc2 tumor cells were co-inoculated with RAW264.7-derived M2 macrophages expressing high levels of TIM3 or with TIM3 knockout (two independent clones). Eight-week-old C57BL/6 mice were subcutaneously implanted with 5 × 10□ KP-Luc2 cells with macrophages suspended in Matrigel Matrix (Corning) into both flanks with four groups: 1. KP-Luc2 cells alone; 2. KP-Luc2 + 5 × 10□ RAW264.7 (TIM3high) (control); 3. KP-Luc2 + 5 × 10□ RAW264.7-TIM3-KO#1; 4. KP-Luc2 + 5 × 10□ RAW264.7-TIM3-KO#2. Tumor volumes were measured using a digital caliper twice weekly beginning on day 5 post-inoculation. At week 3, mice were euthanized, and tumors were excised, photographed, and weighed.

##### In Vivo Immunotherapy: aTIM-3/aPD-1 and aTIM-3/aPD-L1 Blockade (see Figure 7A-7D; Supplemental Figure 7D-7E)

For immune-checkpoint inhibition studies, C57BL/6 mice were subcutaneously injected with 5 × 10□ KP-Luc2 cells bilaterally. Mice received anti-TIM3 (aTIM-3) and/or anti-PD1 (aPD-1) neutralizing antibodies (10 mg/kg each, i.v. route, twice weekly). Tumor size was monitored twice weekly, and tumors were collected on day 17 for imaging and weight quantification (referred to Figure 7A-7D). A similar designed experiment was performed using aTIM-3, aPD-L1 and their combination using the same dosing and treatment schedule.

##### Macrophage Depletion Using Clodronate Liposomes (see Supplemental Figure 7A-7C)

To assess the contribution of TIM3□ macrophages to tumor growth, macrophages were depleted using clodronate liposomes. C57BL/6 mice were subcutaneously inoculated bilaterally with KP-Luc2 cells. On day 5 post-tumor implantation, mice received either control liposomes or clodronate liposomes via intravenous retro-orbital injection. Two days later, aTIM-3 antibody was administered intravenously. This combination treatment was repeated weekly for a total of two cycles.

##### Immunotherapy + Chemotherapy in Peritoneal Metastatic (PM) Model (see Figure 7F-7I and Supplemental 7G-7I)

To model peritoneal metastasis and mimic the mitomycin C (MMC) HIPEC treatment paradigm in the clinic, mice were intraperitoneally injected with KP-Luc2 cells, followed by immunotherapy and/or chemotherapy as outlined in Figure 7F. Mice were randomized into six treatment groups: (1) isotype IgG control; (2) αTIM-3; (3) αTIM-3 + MMC (1 mg/kg; Sigma); (4) MMC alone; (5) αPD-1; and (6) αTIM-3 + αPD-1 + MMC. Treatments were administered twice weekly for three weeks. Tumor burden was monitored weekly by bioluminescence imaging. At the study endpoint, mice were euthanized, and peritoneal tumor nodules were collected, counted, and weighed. Tumor tissues were either fixed in 4% formalin for histopathological analysis or enzymatically dissociated into single-cell suspensions for flow cytometric immune profiling using antibodies listed in Supplemental Table 3.

#### Statistical Analysis

Statistical analyses of flow cytometry, immune response, and selected scRNA-seq datasets were performed using an unpaired, two-tailed *t* test, one-way ANOVA with Tukey’s multiple-comparison test, or Fisher’s exact test in GraphPad Prism (GraphPad Software). TCGA datasets were obtained from cBioPortal, KMplot, and GEPIA (https://gepia.cancer-pku.cn). Kaplan–Meier survival curves were generated, and statistical differences were assessed using the log-rank (Mantel–Cox) test. Data were confirmed to meet the assumptions of each statistical test; when variances were unequal (as determined by an F test), Welch’s correction was applied. A *P* value < 0.05 was considered statistically significant. Error bars represent the standard error of the mean (SEM). When multiple visual fields were averaged to generate a single value per animal, these animal-level values were then averaged to obtain the group mean shown in each graph.

## Results

### TIM3 and its ligands are increased in metastatic cells/tissues compared to primary GC tissues and associated with poor survival

Our prior bulk RNAseq study of 44 GCPM specimens revealed much higher expression levels of TIM3 (*HAVCR2*) than PD-1, PD-L1, CTLA4, and TIGIT^18^. We also noted that expression of galectin-9 (Gal-9; *LGALS9*), one of TIM3’s major ligands was very high in the same samples, with higher TIM3 and Gal-9 expression in GCPM (particularly mesenchymal type GC), in association with therapy resistance^18^. Further analysis of Bulk RNAseq from GCPM revealed that TIM3 is highly associated with notable tumor immune suppressive markers including CD276, CD163, CD11b(ITGAM), CD68, and LAG3 etc. especially in immune high and tumor cell percentage low clusters (Supplemental Figure 1A). To further confirm these findings, we examined TCGA GC dataset, finding that TIM3 and its ligands Gal-9, CEACAM1, and HMGB1 are significantly increased in primary GC tissues (n=408) compared to normal tissues (n=211) (Figure 1A and Supplemental 1B&1C). Significantly increased expression of TIM3 in GC tumor tissues compared with normal tissues were further validated in two independent GC cohort (GES33335 and GSE146996) respectively (Figure 1B). Moreover, TIM3 expression was much higher in peritoneal metastatic tumors than that of primary tumor tissues with or without PM from a recent published dataset^19^(p=0.0041, p=5.9e-05) (Figure 1C). To further validate TIM3 expression in GC metastatic cells and tissues compared with primary tumors, we examined TIM3 expression using immunofluorescent staining (co-IF) in primary and gastric cancer peritoneal metastases (GCPM) specimens and found TIM3 is highly enriched in GCPM samples compared to primary tumors (Figure 1D). Our prior study has identified that SOX9, a tumor marker in GCPM, plays a critical role in GCPM^20^, while TIM3 is exclusively expressed in SOX9 negative non tumor cells in GCPM samples (Figure 1D, lower panel). Furthermore, Boxplot comparisons of TIM3 (*HAVCR2*) expression across cell types between primary tumors and peritoneal metastasis from a single-cell RNAseq of independent GC cohort^21^ further revealed that TIM3 is significantly enriched in macrophages of peritoneal metastases (PMs) compared to primary GC but not in other cell types (Figure 1E). The increased expression of TIM3 along GC progression was further confirmed by co-IF in normal, additional primary tumor and GCPM cases (Supplemental Figure 2A). To further explore their clinical relevance, we found higher TIM3 expression and that of its ligands Gal-9, CEACAM1 and HMGB1 in significant association with poor survival respectively (https://kmplot.com) in GC tissues (Figure 1F&Supplemental Figure 2B). This suggests that TIM3 along with its ligands, while expressed in primary GC tissues, is much higher in GCPM, likely plays an important role in GC progression and metastases.

**Figure 1.**
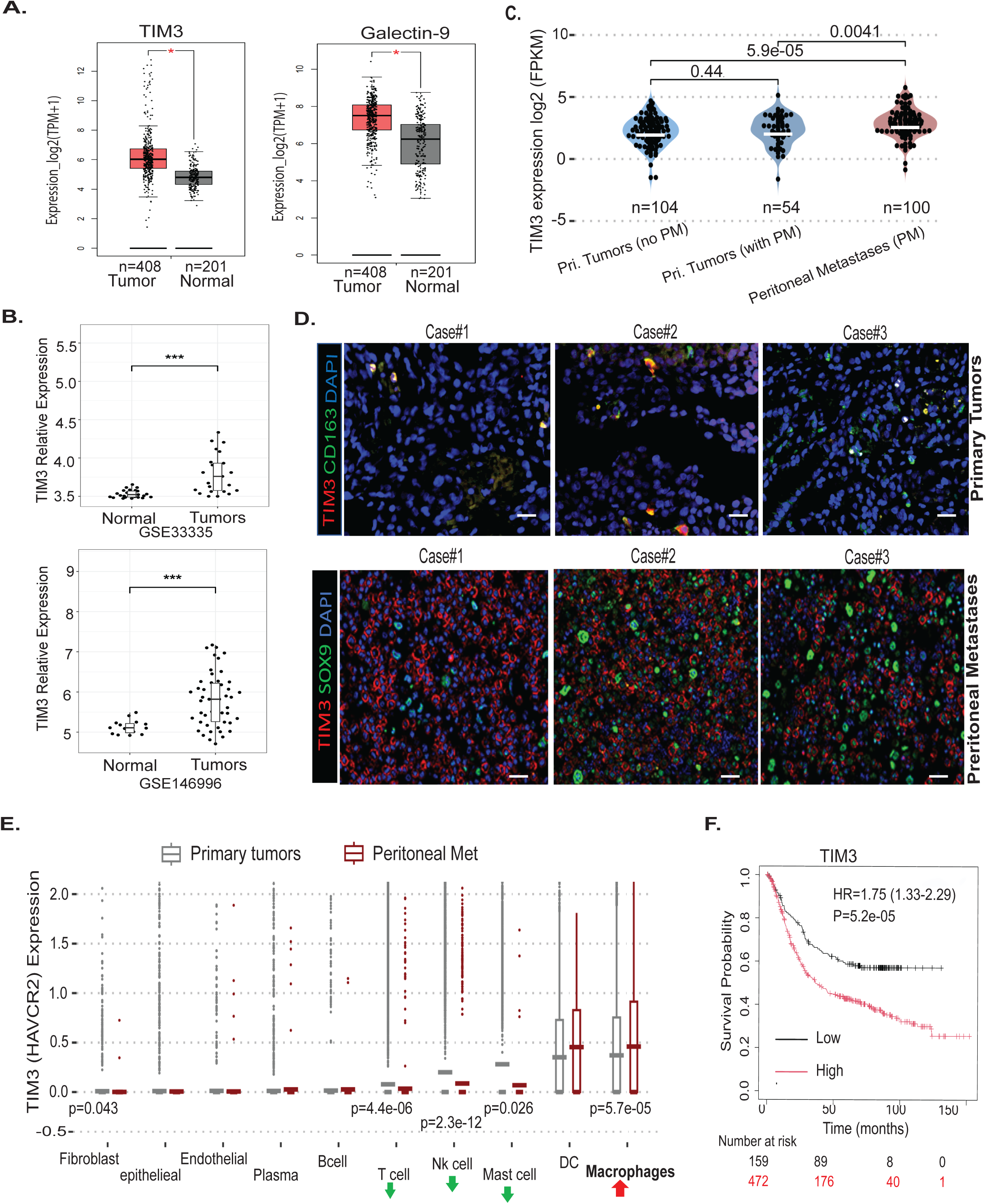
TIM3 and its ligands are increased in metastatic cells/tissues compared to primary GC tissues and associated with poor survival. **A.** Increased expression of TIM3 and its ligand Galectin-9 (Gal-9) in GC tumor tissues (n=408) compared to normal tissues (n=201), analyzed using the GEPIA database (http://gepia.cancer-pku.cn). *p<0.05. **B.** Expression of TIM3 is significantly higher in GC tumor tissues than normal in two independent cohorts (GSE33335 and GES146996) (\*\*\**P*<0.001); **C.** Violin plot comparing TIM3 (*HAVCR2*) log2(FPKM) gene expression across samples retrieved from GC patients with and without peritoneal metastasis (PM) from a previously published dataset described ^19^. Statistical comparisons were undertaken using a two-sided T test. **D.** Representative co-immunofluorescence staining of TIM3/CD163 and TIM3/SOX9 in primary GC tumors and GC peritoneal metastases (GCPMs). Scale bar, 20 μm. **E.** Boxplot comparisons of TIM3 (*HAVCR2*) expression across cell types between primary tumors and peritoneal metastasis from a recent published GC cohort^21^. F. Kaplan–Meier analysis showing the association between TIM3 expression and overall survival in >600 advanced GC patients from the TCGA database (HR = 1.75, P = 5.2 × 10; https://kmplot.com).

### TIM3 is dominantly enriched in tumor associated M2 macrophages in GCPM samples

Next, we determined which cell types, in GCPM samples, primarily express TIM3. Our scRNAseq data analyses, in all cell types from 20 GCPM revealed that expression of TIM3 is primarily in macrophages among different cell population of GCPM samples in violin plot (Figure 2A), and TIM3 expression coexist with notable macrophage markers CD163 and GSF1R in GCPM samples (Supplemental Figure 3A). Next, we measured expression of TIM3 and its ligands Gal-9 and CEACAM1 in 54 GCPM specimens using multi-flow cytometry analysis on different cell types using notable cell type markers (epithelial tumor cells-EpCAM, immune cells-CD45, M2 macrophages-CD163, and CD4 and CD8 for T cells). As indicated in Figure 2B, we confirmed that TIM3 is highly enriched in CD163+ macrophages, while it is minimal in EpCAM+ tumor cells and CD3, CD4, and CD8 T cells. Likewise, in addition to express in CD163+ macrophages, Gal-9 was widely expressed in tumor cells and other immune cells, while CEACAM1 was primarily in tumor cell clusters (Figure 2B), consistent with our scRNAseq data (Supplemental Figure 3B). Expression details of TIM3, its ligands Gal-9, CEACAM1 and PD1 and PD-L1 in all different cell populations in the 54 GCPM specimens are shown in Supplemental Table 1. CyTOF data analyses of 6 GCPM cases also revealed that TIM3 coexists with the macrophage marker CD68 (Figure 2C). A significantly high correlation between TIM3 and CD163 and its ligand Gal9 was also confirmed in our scRNAseq of 20 GCPM cases (Figure 2D). Further TCGA analysis found a high correlation between TIM3 (*HAVCR2*) and the notable M2 macrophage markers CD163 (R=0.66,p-value =0) and CSF1R (R=0.79, p-value=0) (Figure 2E), CD68 (R=0.71, p-value=0) and PD-L1 (R=0.27, p-value=3.9e-08) (Supplemental Figure 3C). The significant high correlation between TIM3 and notable M2 macrophage marker CD163 was further validated in two GC cohorts (GSE33335 and GSE146996) and a metastatic GC cohort (GSE237876) (Supplemental Figure 3D). Furthermore, the higher expression of TIM3 in macrophages was further validated from a recent published independent single cell RNA-seq profiling of peritoneal tumor samples^21^ in UMAP plot (Supplemental Figure 3E). Finally, using co-immunofluorescent staining of TIM3 and CD163, we further validated that both TIM3 and CD163 are highly co-localized in M2 macrophages in more than 30 GCPM specimens (Figure 2F). All these data indicate that TIM3 is primarily expressed in tumor associated M2 macrophages in GCPM specimen.

**Figure 2.**
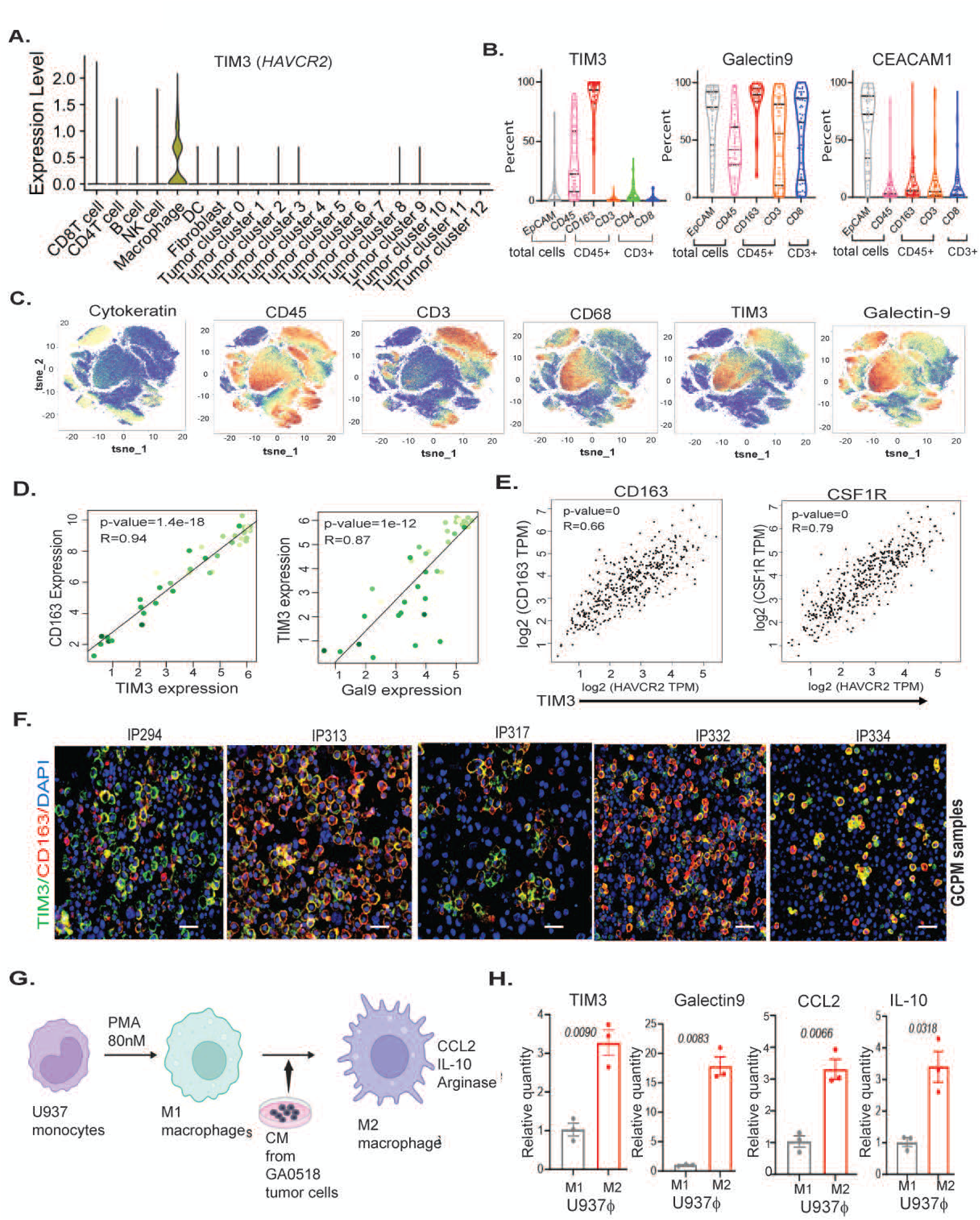
TIM3 is dominantly enriched in tumor-associated M2 macrophages in GCPM samples and induced by tumor-conditioned medium. **A.** Violin plot showing TIM3 expression across all cell types from scRNA-seq analysis of unfractionated live cells from 20 GCPM samples. **B**. Multi-parameter flow cytometric analysis of TIM3 and its ligands Gal-9 and CEACAM1 across different cell types in 54 GCPM samples (see Supplemental Table 1). **C**. CyTOF analysis of TIM3, Gal-9, and immune markers in six GCPM samples, using cytokeratin, CD45, CD34, and CD68 to define epithelial and immune populations, as described in Methods. **D**. Strong positive correlations between TIM3 and CD163 (R = 0.94, P = 1.4 × 10□¹□) and between TIM3 and Gal-9 (R = 0.87, P = 1 × 10□¹²), validated by scRNA-seq analysis. **E**. Positive correlations between TIM3 and M2 macrophage markers CD163 and CSF1R in GC tissues, analyzed using the GEPIA database. **F**. Representative co-immunofluorescence staining of TIM3 and CD163 in five GCPM samples. Scale bar, 20 μm. **G**. Schematic illustrating macrophage polarization from U937 monocytes to an M1 state followed by repolarization to an M2 state using GA0518 tumor cell–conditioned medium. **H**. qRT-PCR analysis of TIM3, Gal-9, and M2-associated markers (CCL2 and IL10) in U937-derived M1 and M2 macrophages. *P* values are indicated.

### Conditioned medium (CM) from tumor cells of human and mouse promoted TIM3 expression and M2 macrophage repolarization

To elucidate if TIM3 increased along macrophage repolarization, using phorbol 12-myristate 13-acetate (PMA, 80nM), we first polarized human U937 monocytes and murine RAW264.7 monocytes into M1 macrophages after 48-hour treatment (Figure 2G). These cells were further polarized into M2 macrophages with tumor cell culture medium (CM) (Figure 2G and Supplemental 4A). As shown in Figure 2H, CM from human GA0518 tumor cells, a tumor cell line derived from GCPM^22^, significantly increased the M2 macrophage markers CCL2, IL-10 as well as increased TIM3 and Gal-9 in human U937 monocytes after repolarization to M2 macrophages. Similarly, CM from KP-Luc2 murine tumor cells significantly increased M2 macrophage markers CCL2, IL-10, and arginase 1 in freshly isolated bone marrow macrophages from mice (mBM Mf) (Supplemental Figure 4A). Interestingly, expression of TIM3 and its ligand Gal-9 were significantly higher in murine M2 macrophages upon tumor cell CM treatment, compared to M1 macrophages (Supplemental Figure 4B). Collectively, these data suggest that TIM3 and its ligand Gal-9 are highly increased in tumor associated M2 macrophages; and induced by tumor cell CM in both human and mice macrophages. Thus, TIM3 might be a critical inhibitory receptor enriched in M2 macrophages to mediate M2 macrophages’ protumor or pro-metastatic functions in GCPM.

### Depletion of TIM3 in macrophages reduced tumor cell invasion, and suppressed tumor growth *in vivo*

To elucidate the functional role of TIM3^+^ tumor associated M2 macrophages in mediating tumor cell malignant behaviors, we first knocked out (KO) TIM3 in both human U937 and mouse RAW264.7 monocyte lines by lentiCRISPR/CAS9 and then repolarized them into M2 macrophages by adding CM from human GA0518 tumor cells or murine KP-Luc2 tumor cells. These M2 macrophages were co-inoculated with GA0518 tumor cells (upper level) as indicated in Figure 3A and then determine their influence on tumor cell malignant behaviors. As a result, we found that co-inoculation of GA0518 tumor and U937 TIM3^high^ control cells significantly increased GA0518 tumor cell invasive capacity compared to GA0518 alone, while TIM3 depletion from human U937 macrophages dramatically reduced tumor cell invasion (Figure 3B). Successful TIM3 KO in U937 cells was confirmed by western blot (left panel of Figure 3B). Similar results were also demonstrated in the mouse system, with significantly fewer invading Kp-Luc2 mouse tumor cells upon co-inoculation with RAW264.7 TIM3 KO in two individual clones compared to the RAW264.7 control (Figure 3C). This was a sturdy observation that both human and mouse M2 TIM3^high^ macrophages greatly increased tumor cell invasion compared to tumor cells alone, while TIM3 KO in M2 macrophages significantly reduced tumor cell invasion.

**Figure 3.**
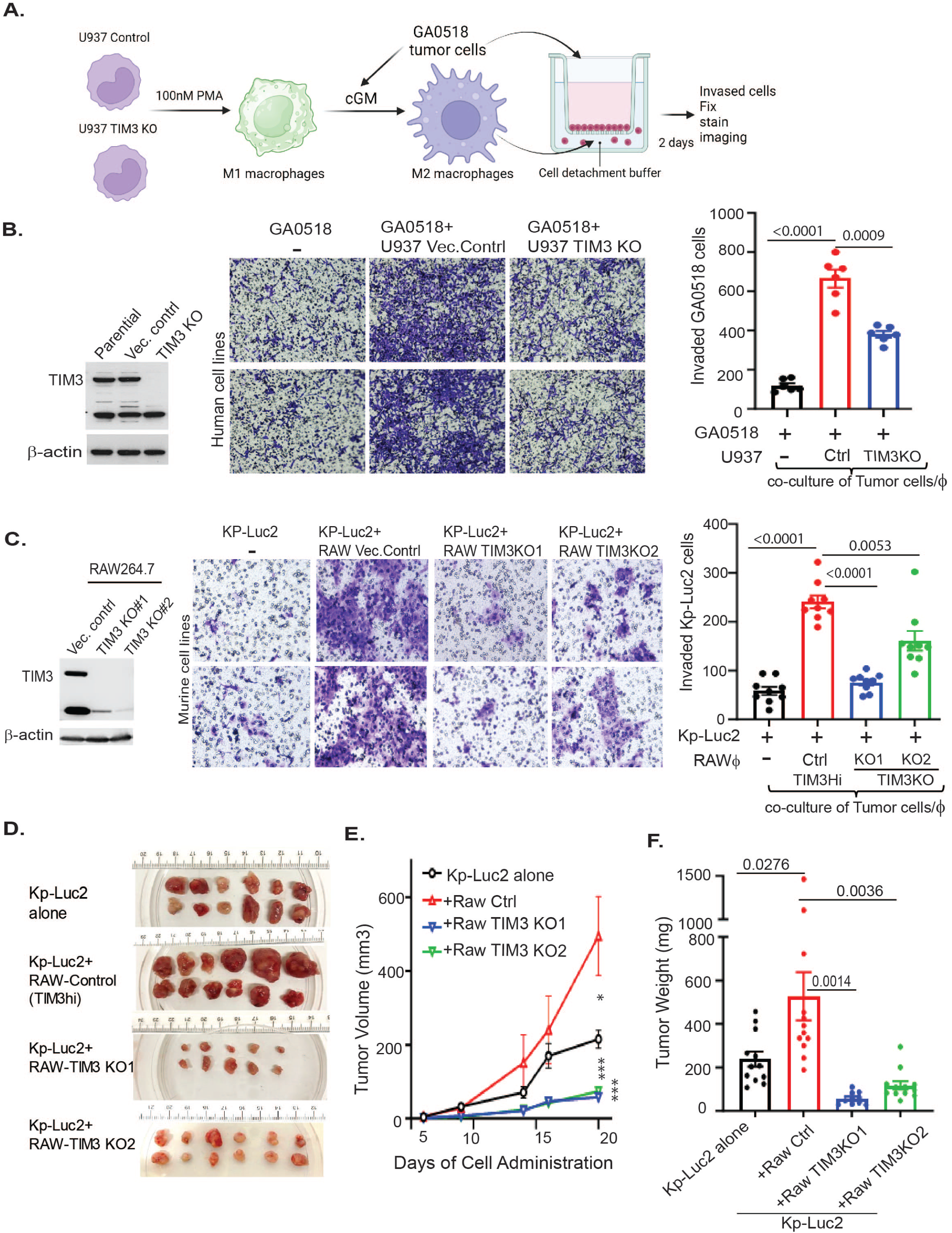
TIM3 depletion in macrophages reduces tumor invasion and suppresses tumor growth in vivo. **A.** Schematic of the in vitro tumor cell invasion assay toward M2 macrophages with or without TIM3 knockout (KO). **B.** Effects of TIM3 KO in human U937 macrophages on GA0518 tumor cell invasion. Left: Western blot validation of TIM3 KO. Middle: representative invasion images. Right: quantification of invaded cells. **C.** Effects of TIM3 KO in murine RAW264.7 macrophages on KP-Luc2 tumor cell invasion, with validation, representative images, and quantification. .**D-F**. Co-inoculation of KP-Luc2 tumor cells with RAW264.7-derived M2 macrophages expressing TIM3 or TIM3 KO. (**D**) Representative tumor images. (**E**) Tumor growth curves and volumes. (**F**) Tumor weights at the experimental endpoint.

Next, we further validated the tumor-promoting role of TIM3+ macrophages *in vivo* in an immune-competent KP-Luc2 syngeneic mouse model. Here, we subcutaneously co-inoculated murine KP-Luc2 GC cells^20^ with murine macrophage RAW264.7 control or TIM3KO cells into the flanks of C57BL/6 mice and observed tumor growth over a one-month period. Strikingly, KP-Luc2 tumors grew faster and larger in tumor size and weight when co-injected with TIM3^high^ M2 macrophage controls compared to the tumor cell alone; while co-injection of two individual TIM3 KO clones with KP-Luc2 tumor cells significantly reduced KP-Luc2 tumor cell growth *in vivo* (Figure 3D-3F). Thus, TIM3^high^ M2 macrophages significantly promote tumor cell invasion and tumor growth in GC.

### Depletion/blockage of TIM3 in macrophages increases human and mouse T cell immunity *in vitro* and *in vivo*

Since it was reported that TIM4+ cavity-resident macrophages impair CD8 T cell proliferation and anti-tumor immunity^23^, we tested whether TIM3 in macrophages might affect T cell functions using an *in vitro* co-culture system. First, human U937 or murine RAW247.6 monocytes with or without TIM3 KO were repolarized into M2 macrophages using PMA and CM from GA0518 or KP-Luc2 tumor cells respectively, and then co-cultured with human PBMCs for 48 hours. The suspended T cells were then collected and analyzed by flow cytometry (Figure 4A). As shown in Figure 4B, the number of CD3 and CD8 T cells significantly increased when co-cultured with U937 TIM3KO cells compared to those in U937 control cells. Consistently, production of interferon-g and perforin in CD3 and CD8 T cells was significantly increased upon co-culture with U937 TIM3 KO cells compared to U937 control (Figure 4B). Similar findings were noted in co-cultured murine PBMCs and RAW264.7 macrophages (Figure 4C). Likewise, TIM3 KO in RAW264.7 macrophages significantly stimulated IFNg and perforin production from CD3 and CD8 T cells respectively that were corresponding to the increase in CD3 and CD8 T cell numbers compared to the control vector group (Figure 4C & Supplemental Figure 5A).

**Figure 4.**
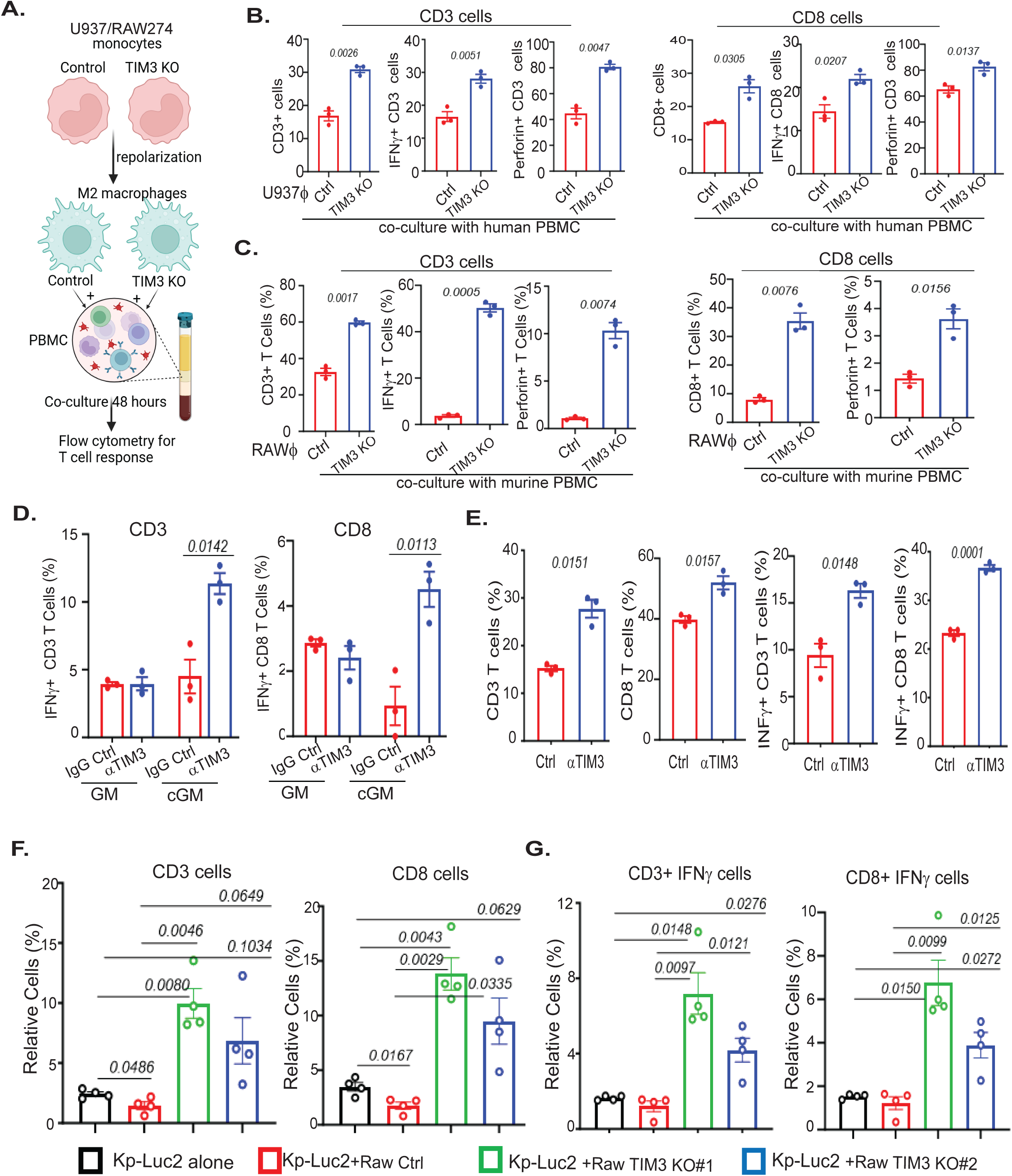
TIM3 depletion or blockade in M2 macrophages enhances T cell responses. **A.** Schematic of in vitro co-culture of human or murine macrophages (control vs TIM3 KO) with PBMCs. **B**, Flow cytometry analysis of CD3□ or CD8□ T cells and intracellular cytokines, interferon-γ (IFNγ) and perforin in PBMCs after co-cultured with U937-TIM3 KO cells compared with U937-Control (Ctrl) cells. **C**, Flow cytometry of CD3□ or CD8□ T cells and intracellular IFNγ and perforin in PBMC co-cultured with murine RAW-TIM3KO macrophages versus RAW-Control. **D.** co-culturing of murine PBMCs with mouse bone-marrow-derived macrophages (mBM-Mø) under tumor cell derived conditioned medium (cGM) vs growth medium (GM), followed by anti-TIM3 antibody treatment. **E**. Flow cytometric quantification of CD3□/CD8□T cells and IFNγ production in CD45□ immune cells from malignant ascites co-cultured with U937 macrophages treated with anti-TIM3 antibody. **F-G.** Flow cytometric analysis of CD3 /CD8 T cell numbers (F) and IFNγ production (G) in tumors from the KP-Luc2 syngeneic model co-inoculated with RAW264.7 macrophages.

To further confirm the effects of macrophagic genetic TIM3 KO on T cell response, we pharmacologically inhibited TIM3 using an anti-TIM3-neutralizing antibody. As shown in Figure 4D, mouse bone marrow-derived monocytes (BMDMs) were used as native macrophages and co-inoculated with mouse PBMCs, the production of interferon g from CD3 and CD8 T cells was significantly higher upon treatment with anti-mouse TIM3-neutralizing antibody in the presence of tumor CM (cGM) than that of normal culture medium (GM) (Figure 4D). Similarly, CD45+ immune cells (containing TIM3+ macrophages and CD3+ T cells) from patients’ malignant ascites treated with anti-TIM3-neutralizing antibody significantly increased the CD3 and CD8 T cell numbers and their production of interferon-g (INFg) from CD3 and CD8 T cells respectively (Figure 4E). Similarly, INFg, GranB, and perforin from CD3+ T cells from another patient (GA071924) was significantly increased upon anti-TIM3 treatment compared to control (Supplemental Figure 5B). These results demonstrate that T cell response was stimulated upon TIM3 blockage by its neutralizing antibody. Furthermore, a tumor cell-killing assay illustrated that T cells (from PBMCs) co-cultured with U937 TIM3KO cells significantly killed GA0518 tumor cells more compared to the U937 vector control group (Supplemental Figure 5C).

Depletion of TIM3 in murine RAW264.7 M2 macrophages significantly reduced KP-Luc2 tumor growth in the syngeneic model (Figures 3D-3F). To elucidate if depletion of TIM3 from macrophages blunts tumor growth due to boosting T cell responses *in vivo*, we analyzed CD3 and CD8 T cell infiltration, and interferon-g production in KP-Luc2 mouse tumors co-inoculated with TIM3^high^ or TIM3 KO RAW264.7 macrophages after repolarization to M2 state. As a result, TIM3 KO in RAW264.7 cells vastly increased CD3 and CD8 T cell numbers in tissues with co-innovation of tumor cells with RAW264.7 TIM3 KO clones compared to control (Figure 4F). Consistently, interferon-g production from CD3 and CD8 cells were significantly increased in tumors with co-injection of RAW264.7 TIM3 KO clones compared to Kp-luc2 alone or RAW264.7 control with TIM3 high (Figure 4G). This suggests that TIM3^+^ tumor associated M2 macrophage suppresses T cell functions in the GC TME.

### TIM3 dictates the secretome of cytokines and chemokines from M2 macrophages

We speculated that cytokines/chemokines from TIM3^+^ tumor associated macrophages might determine macrophage functions on tumor cells and T cell immunity. After further confirming efficient KO of TIM3 in U937 M2 macrophages using co-IF and Western blot (Figure 5A&5B), we first determined changes of the three notable M2 macrophage markers CCL2, IL-10, and Arginase 1. As expected, depletion of TIM3 from human U937 macrophages dramatically reduced CCL2 and IL10 levels compared to vector control and parental cells (Figure 5C). Similarly, CCL2, Arginase 1 and IL10 were significantly suppressed upon TIM3 depletion from murine RAW267.4 cells (Figure 5D). To comprehensively determine the secretome (cytokines/chemokines) influenced by TIM3 in macrophages that might mediate tumor-promoting and immune-suppressive functions, we performed cytokine arrays on CM from U937 vector control (VB) and U937 TIM3 KO. As indicated in Figure 5E, among 28 cytokines detected, 7 cytokines/chemokines CCL1, CCL4, CCL5, CCL7, IL-1b, IL-6, and CCL20 (MIP-3a) were significantly reduced upon macrophagic TIM3 depletion that was further validated by qRT-PCR in U937 cells (Figure 5F). These data support macrophagic TIM3 strongly dictates the production of key cytokines/chemokines that might mediate the interactions among M2 macrophages, tumor cells, and T cell function in the TME, while also exerting immune-suppressive functions by TIM3^+^ macrophages to promote GCPM.

**Figure 5.**
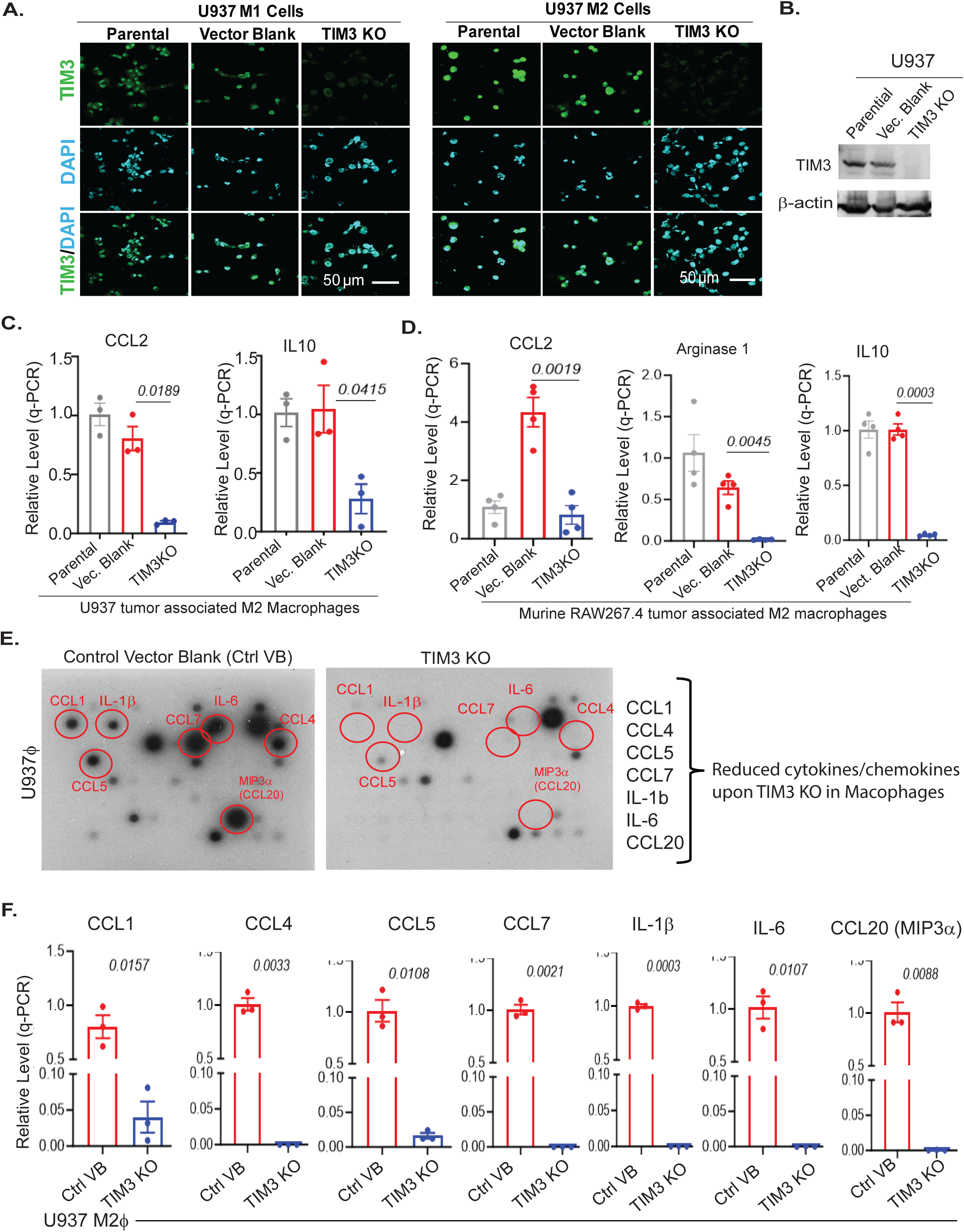
TIM3 in macrophages dictate secretome of cytokines/chemokines from M2 macrophages. **A.** Immunofluorescence staining of TIM3 in U937-derived M1 and M2 macrophages with or without TIM3 KO. **B.** Western blot confirmation of TIM3 KO in U937 macrophages. **C.** qRT-PCR analysis of CCL2 and IL10 expression in U937 M2 macrophages with or without TIM3 KO. **D.** qRT-PCR analysis of murine M2 macrophage markers (CCL2, Arg1, and IL10) in RAW264.7 macrophages with or without TIM3 KO. **E**, Cytokine array profiling of secreted factors from U937 TIM3 KO versus control cells using the RayBio C-Series Human Cytokine Antibody Array. **F**, Validation of seven selected cytokines/chemokines by qRT-PCR. *P* values are indicated.

### TIM3^+^ tumor associated M2 macrophages promote tumor invasion and suppress T cell function partially via the p90RSK1/2/CCL20 axis

To elucidate how TIM3□ tumor associated M2 macrophages mediate metastatic and immunosuppressive functions in GC tumor microenvironment (TME), we hypothesized that key cytokines released by TIM3□ macrophages—identified by cytokine array analysis (Figure 5E)—serve as critical effectors. Among these candidates, C-C motif chemokine ligand 20 (CCL20, also known as MIP-3α) was the most abundant cytokine in GCPM specimens, detected in both tumor cell clusters and macrophages by scRNA-seq (Figure 6A and Supplemental Figure 6A). CCL20 is well documented to be upregulated across multiple tumor types, including GC. This chemokine recruits CCR6□ cells—encompassing immune populations such as Tregs and Th17 cells as well as tumor cells—thereby promoting cancer cell migration and invasion, and suppressing T-cell function through CCR6 signaling^24, 25^. However, direct evidence for a functional role of the CCL20/CCR6 axis in GC remains limited. Using TCGA data, we observed that both CCL20 and its receptor CCR6 are markedly elevated in GC (STAD) tumor tissues relative to normal controls (Supplemental Figure 6B). Notably, among TIM3-regulated cytokines in macrophages, CCL20 showed the significant increase in tumors vs normal compared with other candidates (e.g., CCL1, CCL4, CCL7) across two independent GC cohorts (Figure 6B and Supplemental Figure 6C). These findings led us to focus on CCL20 as a key mediator of TIM3-dependent macrophage function in GCPM in the subsequent studies. To assess CCL20 expression and production from different states of macrophages, we repolarized U937 monocytes to M1 and M2 states and measured CCL20 expression levels in the macrophage using qRT-PCR and ELISA. As expected, CCL20 mRNA and protein secretion were much higher in M2 macrophages than that in M1 macrophages (Figure 6C). Interestingly, TIM3 knockout (KO) in both human (U937) and mouse (RAW274.6) macrophages dramatically reduced CCL20 secretion by ELISA even when repolarized to the M2 state (Figure 6D) indicating that TIM3 in macrophages dictates CCL20 production and secretion.

**Figure 6.**
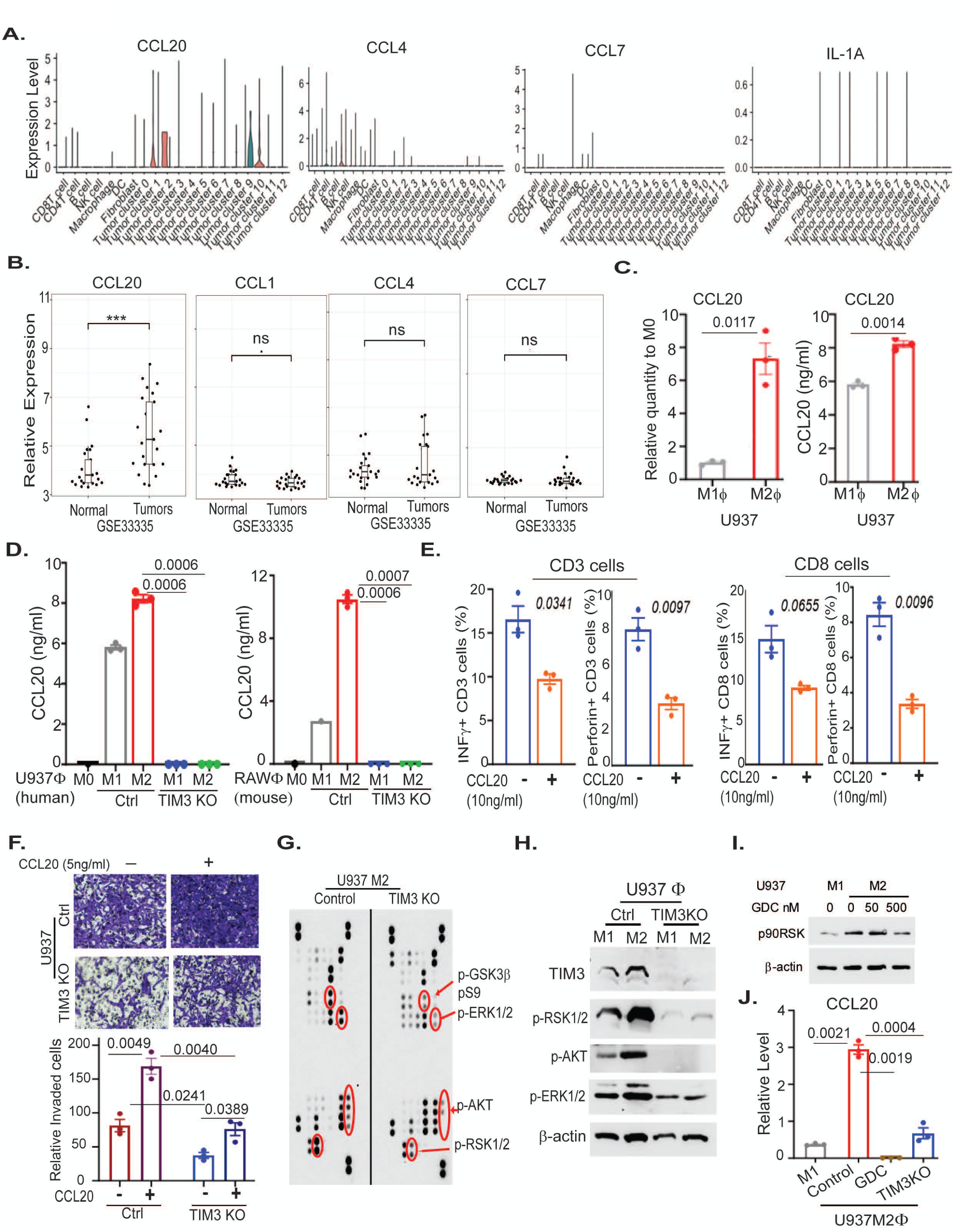
TIM3D M2 macrophages promote tumor invasion and suppress T cell function via the p90RSK1/2–CCL20 axis. **A.** Violin plots showing CCL20, CCL4, CCL7, and IL1A expression across cell types in 20 GCPM samples by scRNA-seq; **B.** Relative expression of CCL20, CCL1, CCL4, and CCL7 in GC tumor versus normal tissues from the GSE33335 cohort. **C.** CCL20 expression in U937-derived M1 and M2 macrophages assessed by qRT-PCR and ELISA. **D.** ELISA measurement of secreted CCL20 from human U937 and murine RAW264.7 macrophages with or without TIM3 KO. **E**, Flow cytometric analysis of IFNγ and perforin production in CD3 and CD8 T cells co-cultured with U937 macrophages in the presence of recombinant CCL20. **F.** Tumor cell invasion toward U937 control or TIM3 KO macrophages in the presence of recombinant CCL20. **G**. Phospho-kinase array analysis comparing U937 TIM3 KO and control macrophages after M2 repolarization. **H**, Western blot validation of TIM3 and selected kinases in U937 macrophages. **I.** Effect of GDC-0994 on phosphorylation of p90RSK1/2 was determined by Western blot analysis. **J.** qRT-PCR analysis of CCL20 expression following GDC-0994 treatment compared with TIM3 KO.

Based on the above observations, we proposed that CCL20 from TIM3^+^ macrophages might mediate TIM3’s tumor-promoting and immune-suppressive functions. Thus, recombinant CCL20 (rCCL20) might retard the T cell immune response. When U937 macrophages were cocultured with PBMCs, we found that rCCL20 significantly reduced interferon g and perforin production from CD3 and CD8 T cells respectively (Figure 6E). Furthermore, while TIM3 KO in U937 cells significantly reduced GA0518 tumor cell invasion, this was dramatically increased by rCCL20 upon co-culture with U937 macrophages with or without TIM3 KO (Figure 6F).

To further elucidate the mechanisms by which TIM3 regulates CCL20 expression in macrophages, a kinase array including 22 phosphorylated kinases was performed on TIM3^+^ M2 macrophages paired with TIM3 KO U937 cells. As indicated in Figure 6G, phosphorylation of four kinases (p90RSK1/2, p-GSK3b, p-ERK1/2, p-AKT) was dramatically reduced upon TIM3 KO in U937 M2 macrophages. We further validated these four kinases by western blot, finding that phosphorylation of p90RSK1/2 and its upstream kinase ERK1/2 was dramatically increased in TIM3^+^ M2 macrophages compared to M1 macrophages, while TIM3 KO dramatically reduced p90RSK1/2 and ERK1/2 activation as reflected by its phosphorylation indicating that ERK1/2/p90RSK1/2 axis is the major kinases regulated by TIM3 on CCL20 expression (Figure 6H). Furthermore, we have demonstrated that p90RSK1/2, an active form of the RSK1/2, increased in TIM3 high M2 macrophages but reduced by ERK1/2 inhibitor GDC-0994 at 500nM (Figure 6I). Most importantly, increased CCL20 in M2 macrophages significantly reduced by GDC-0994 that showed the similar effects to that of TIM3 KO (Figure 6J) suggesting TIM3 in M2 macrophages activates ERK1/2/p90RSK1/2 that increase CCL20.

Moreover, TIM3 high M2 macrophages stimulate tumor cell invasion *in vitro* and tumor growth *in vivo* (Figure 3D-3F), to determine the activated signaling of tumor cells induced by TIM3^high^ M2 macrophages, we cultured GA0518 tumor cells with CM from TIM3^high^ or TIM3^KO^ of U937 cells. As shown in Supplemental Figure 6D, phosphorylation of ERK1/2 and p90RSK1/2 was dramatically increased in GA0518 cells under CM of TIM3^high^ U937 cells, but reduced under CM of TIM3^KO^ U937 cells, while ERK1/2 inhibitor GDC-0994 strikingly reduced phosphorylation of ERK1/2 and p90RSK1/2 in GA0518 tumor cells upon co-cultured with U937 TIM3^high^ cells which was further validated by a specific p90RSK1/2 inhibitor BI-D1870. To elucidate the CCL20 from TIM3^high^ macrophages can further activate tumor cell p90RSK1/2 activation through paracrine pattern, under the same co-culture condition, we add rCCL20 in GA0518 tumor cells co-cultured CM from U937 cells with or without TIM3 KO. As a result, we found that CM from TIM3^high^ U937 cells can increased phosphorylation of p90RSK1/2, while CM from TIM3^KO^ U937 cells dramatically reduced activation of p90RSK1/2. Interestingly, rCCL20 further increased phosphorylation of p90RSK1/2 in GA0518 tumor cells stimulated by U937 TIM3^high^ cells; and GDC-0994 can reduce the phosphorylation of p90RSK1/2 induced by rCCL20 (Supplemental Figure 6E). Altogether, these data suggest that TIM3^high^ macrophages affect tumor cell growth through paracrine pattern via CCL20/ERK1/2/p90RSK1/2 signaling.

### TIM3 blockage restricts tumor growth depending on macrophages *in vivo*

Using our syngeneic mouse model, we subcutaneously inoculated mouse Kp-luc2 GC cells into the flanks of C57BL/6 mice. Then, macrophages were depleted by i.v administrating clodronate liposome (used as depletion of macrophages) or control-liposomes, followed by a TIM3-neutralizing antibody, 6 days post-injection. As results, the administration of TIM3 antibody or clodronate liposome macrophage-depletion group displayed profoundly smaller tumor sizes (Supplemental Figure 7A&7B), reduced tumor weights (Supplemental Figure 7C) and slower tumor growth rates than the control group (Supplemental Figure 7C). However, a TIM3-neutralizing antibody did not further reduce tumor size upon macrophage depletion with clodronate-liposomes, indicating that the effects of TIM3 blockage depend on the presence of macrophages *in vivo*.

### TIM3 blockade sensitizes ICB immunotherapy and MMC chemotherapy *in vivo*

Immunotherapies targeting tumor associated macrophages (TAMs) have now advanced to the clinic including disruption of macrophage expansion and differentiation by blocking CSF1R^26^, and we identified that highly enriched TIM3 in tumor associated M2 macrophages in advanced GC mediates immunosuppressive functions in the GCPM TME. To test the antitumor effects of TIM3 blockage in combination with ICB immune therapy i.e.aPD1 or aPDL-1 in an immune-competent mouse model, we assessed a syngeneic murine KP-Luc2 cell line generated from a genetic mouse model of GC^27^. First, we subcutaneously injected KP-Luc2 murine tumor cells, then treated the mice with either anti-TIM3-neutralizing antibody, anti-PD1 antibody that is commonly used in the clinic or their combination for 3 weeks (Figure 7A). As a result, while either treatment alone significantly reduced tumors, combined TIM3 and PD-1 blockage yielded better anti-tumor effects than either treatment alone (Figures 7B-7D). Additionally, the combination treatment significantly increased CD8 T cell infiltration, and decreased expression of the proliferation marker KI67 (Figure 7E). Since we found that PD-L1, the ligand of PD-1, is also highly increased in M2 macrophages in association with TIM3 expression (Supplemental Figure3C, right), we also assessed dual inhibition of TIM3 and PD-L1, in GC growth and metastasis in the KP-Luc2 syngeneic model. Similarly, we found that combining anti-TIM3 and anti-PD-L1 yielded much better anti-tumor activity than either treatment alone (Supplemental 7D-7F). We have an ongoing HIPEC clinical trial using chemotherapy MMC in GI patients with PM. To be consistent with this trial and determine a better therapeutic strategy for patients with PM, we preclinically evaluate the efficacy of combination of TIM3 blockage with MMC chemotherapy with or without anti-PD1 immunotherapy. Thus, we generated an intraperitoneal metastatic model in the immune compromised model by intraperitoneal injection of KP-Luc2 murine tumor cells (Figure 7F), which is a common model for PM that has similar TMEs to that of metastatic human GC^28^. After injecting 5×10^5^ KP-Luc2 cells intraperitoneally 1 week, we treated mice with either anti-TIM3 antibody, MMC, anti-PD1, the combination of anti-TIM3 and MMC and the triple combination of anti-TIM3, MMC and anti-PD1 for three weeks. We found that combination treatment of anti-TIM3 and MMC significantly reduced PM tumor burden by bioluminescence and tumor weight (Figures 7G&7H), while the triple combination of anti-TIM3, MMC and anti-PD1 had further reduced tumor burden and PM volume (Supplemental Figure 7G&7H). More interestingly, in this PM model, we found that IFN-g production from both CD3+ and CD8+ T cells was significantly increased in the combination treatment group compared with either treatment alone in tumor tissues analyzed (Figure 7I). Further, the infiltration of CD8 T cells in the triple combination group was significantly increased in tumor tissues analyzed (Supplemental Figure 7I) compared with control group, whereas other treatment groups showed modest increases. Thus, these data provide a strong rationale for combination of MMC chemotherapy and TIM3 blockage under ICB immune therapy in advanced GC patients with PM.

**Figure 7.**
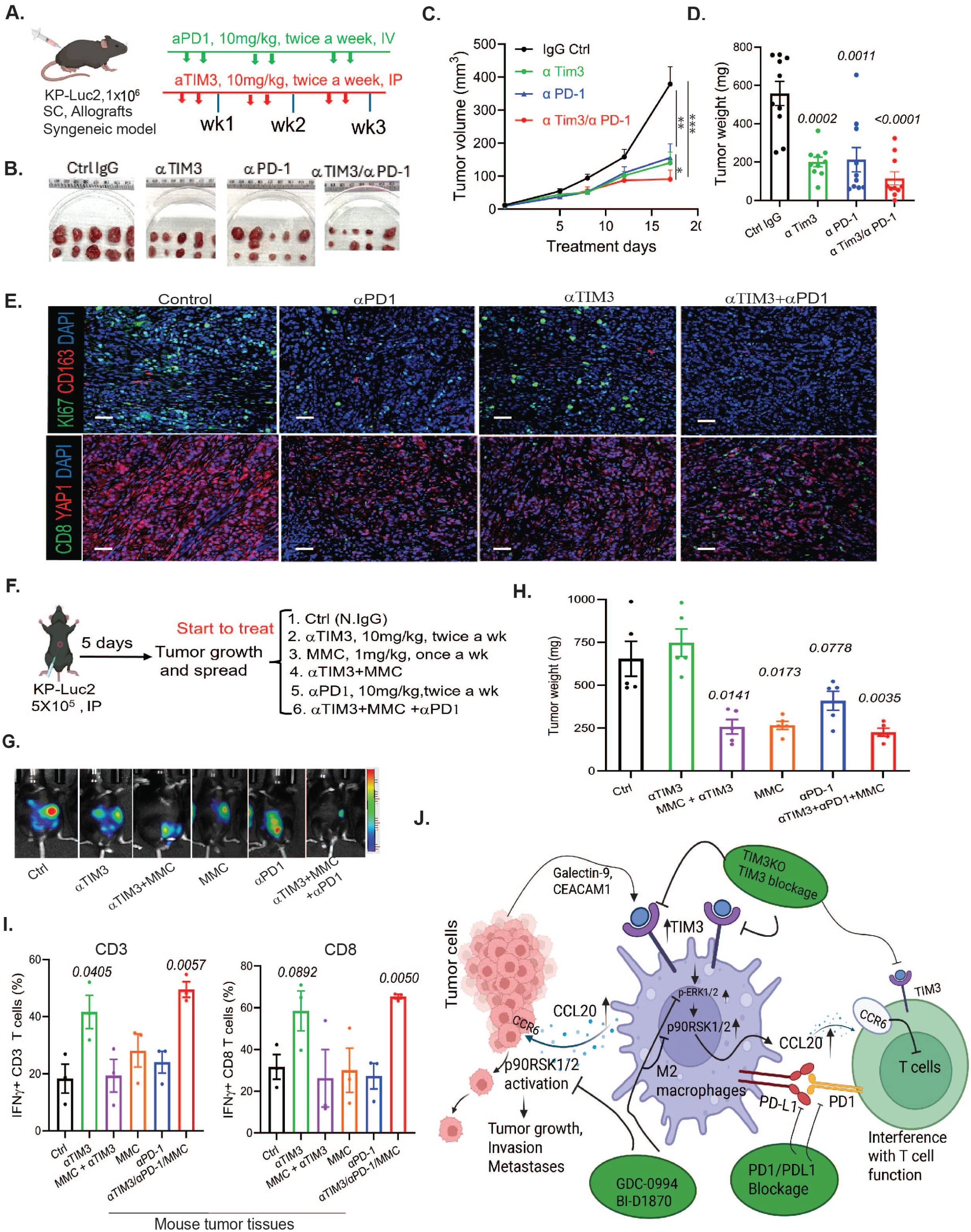
TIM3 blockade, alone or combined with anti–PD-1 or chemotherapy, suppresses tumor growth and metastasis in vivo. **A.** Schematic of the KP-Luc2 subcutaneous tumor model treated with anti-TIM3, anti–PD-1, or combination therapy. **B.** images of tumors from the four treatment groups. **C.** Tumor volume and growth curves over the course of treatment. P value shown at the last measument. *p<0.05, **p<0.01, ***p<0.0001. **D**, Tumor weight (mg) was measured at the experimental endpoint. P value shown in the graph. **E.** Representative co-immunofluorescent staining images of tumor tissues among four group stained with anti-CD163 (red)/Ki67 (green) and YAP1 (red)/CD8 (green). **F.** Schematic illustration of the *in vivo* intraperitoneal (IP) treatment protocol with TIM3/PD1 antibodies combined with mitomycin C (MMC). **G.** Representative bioluminescent images of Kp-Luc2 tumors in six treatment groups were shown. **H.** Average tumor burden/weights was plotted for each treatment group at end of the experiment. P value was shown in the graph. **I.** IFNγ production in CD3 and CD8 T cells from tumor tissues of six treatment group was assessed by flow cytometry. **J.** Schematic diagram illustrating TIM3 in tumor associated macrophages mediated the interactions among macrophages, tumor cells and T cells through p90RSK1/2/CCL20 axis that promote tumor cell invasion, growth and interfere T cell functions, thus supporting the potential novel combination therapies in GCPMs.

## Discussion

GC patients with PM (GCPM) have a very short survival, and overwhelming symptoms and lack effective targeted and immune therapies. The treatment options for advanced GCPM are very limited and its management is highly unsatisfactory^29,30^. Thus, novel immune and target therapeutic strategies for GCPM are urgently needed. In this study, using multiple reverse translational approaches, we have identified that TIM3 is highly enriched in tumor associated M2 macrophages in GCPM, and plays a critical role in promoting tumor cell malignant behaviors, metastases and suppress T cell immune response through remodeling TME. Using cytokine arrays, we found that TIM3^+^ M2 macrophages dictate the secretome of M2 macrophages in TME, and depletion of TIM3 eliminated the productions of notable cytokines that promote metastases; Further analysis revealed that TIM3 in macrophages promote the production of CCL20, and both CCL20 and its receptor CCR6 are highly increased in GC tumors compared to normal tissues and associated with poor survival. Recombinant CCL20 can restore the TIM3 KO function in macrophages on tumor cells and T cells. Further, by kinase array, we identify the p90RSK1/2/CCL20 axis as a critical mediator of the protumor activity of TIM3^+^ macrophages. Finally, we demonstrated that anti-tumor effects of TIM3 blockage depends on macrophages; and abrogation of TIM3 enhances the efficacy of ICB (anti-PD1/anti-PDL1) or mitomycin C (MMC) chemotherapy in preclinical mouse model point to the novel therapeutic strategies for GCPM which can be rapidly translated to the clinic.

It is increasingly appreciated that cancer invasion and metastases result from dynamic crosstalk among tumor cells, macrophages and infiltrating immune cell populations in the TME that rely on a series of bidirectional cellular communications such as ligand–receptor interactions and secreted factors^31^. Tumor associated microphages (TAMs), the most prevalent cell type in the GCPM TME, are crucial in shaping and sustaining a pro-tumorigenic TME by suppressing inflammatory immune responses, facilitating angiogenesis, and fostering a profibrotic microenvironment. These TAM activities contribute to tumor progression, therapy resistance, and tumor dissemination to distant sites ^32, 33,32, 34^.

Our recent profiling of advanced GC samples revealed that CD163+ M2 macrophages are abundant in PM^18, 35^. However, the dominant factors from CD163+ M2 macrophages that mediate immune evasion are still elusive. The key finding of this study is that we showed that TIM3 represents a dominant functional factor in CD163+ M2 macrophages that dictate the secretome from M2 macrophages that potentially mediate metastatic niche and tumor immunosuppressive TME. Cytokine array revealed that KO TIM3 in macrophages significantly reduced the inflammatory cytokines, boost the CD8+ T cell responses either in PBMCs or CD45+ immune cells from malignant ascites and suppress tumor cell invasion. Most importantly, in co-inoculated tumor cells and macrophages *in vivo*, TIM3 KO in murine RAW macrophages significantly reduced tumor growth in the KP-Luc2 syngeneic model (Figures 3D-3F). All these data indicate that TIM3^+^ M2 macrophages play critical roles in promoting tumor growth in GC by remodeling the TME in an immunosuppressive manner.

Mechanistically, we found that macrophage TIM3 promotes production of CCL20, and that both CCL20 and its receptor CCR6 are highly upregulated in GC tumors compared with normal tissues and are associated with poor survival. Recombinant CCL20 restored the pro-tumor effects of TIM3-deficient macrophages on tumor cells and T cells, indicating that the oncogenic activity of TIM3 in macrophages is at least partially mediated through a paracrine CCL20/CCR6 signaling axis. Consistently, in melanoma, tumor cell-expressed CCR6 was found to signal via tumor-associated macrophage (TAM)-derived CCL20, which associated with poor survival and release of the protumoral cytokines TNF and VEGF-A^36^. Finally, we demonstrated that anti-tumor effects of TIM3 blockage depend on macrophages, with TIM3 abrogation enhancing the efficacy of immune checkpoint blockade (anti-PD1/anti-PD-L1) or chemotherapy (e.g., mitomycin C) in preclinical mouse models, pointing to novel GCPM therapeutic strategies that can be quickly translated to the clinic.

Using kinase array, we further identified upregulation/phosphorylation of the kinase p90RSK1/2 (ribosomal protein S6 kinase A1) in TIM3-high macrophages, and its downregulation in TIM3-KO macrophages. This specific kinase is downstream of ERK1/2^37^, which was also inhibited by TIM3 KO. Interestingly, the TIM3 ligand Gal9 is regulated by the estrogen receptor (ER), acting downstream of p90RSK1/2 and ERK1/2, in endometrial cells^38^; thus, it would be interesting to further dissect p90RSK1/2 signaling on regulation of CCL2/CCR6 axis mediated by TIM3 in GCPM. Such studies might also provide insight into our observed effects of TIM KO on cancer cell invasion.

Although our data strongly support TIM3 as a therapeutic target in GCPM, several limitations should be noted. First, despite robust activity in a syngeneic mouse model, additional validation using a Havcr2 (TIM3) knockout mouse model would further strengthen these findings. In addition, evaluation in humanized mouse models would better assess the impact on the human immune system and support the feasibility of translation into human clinical trials.

In summary, we show that TIM3 is highly enriched in tumor associated M2 macrophages in GCPM and plays a critical role in promoting GC progression and suppress T cell functions, thereby contributing to peritoneal metastasis. These findings establish TIM3 as a promising therapeutic target in tumor associated M2 macrophages, in addition to its known role in T cells, representing an immune checkpoint pathway that is similar to but distinct from PD1 and CTLA4 within the TME. Thus, targeting TIM3—particularly in combination with anti-PD1 therapy or MMC chemotherapy—may offer an effective strategy against several devastating malignancies, including GCPM.

## Author contributions

All authors are responsible for the overall content as the guarantors and all authors revised and approved the final version of the manuscript. Conception and, design: SS; Draft the manuscript: SS, XY, JSZ, CB; development of methodology, acquisition of data (performed experiments, analysed data): XY,YF, JSZ, YTZ, DA, MG, AP,JZ, JJ, YH, JM, MT, SD, LW, JS, PT, RS, FS, GG, JAA, SS; acquired and managed patients, provided facilities, resources, PDXs and other support: JAA, JZ,YH, JM,MT, JS, PT, RS, FS, GG, SS; Analysis and interpretation of data: SS, XY, FY, AP, JAA.

## Consent to Publish declaration

All authors have reviewed the manuscript and consent to its publication.

## Consent to Participate declaration

not applicable

## Data availability statement

Data are available in a public, open access repository.

## Patients and Ethics Statement

Ethical approval for this study was obtained from the Institutional Review Board of The University of Texas MD Anderson Cancer Center (LAB01-543) and the Cooper-Coriell Cancer Biobank (IRB CCCB0001), Camden Cancer Research Center. All patients who volunteered to provide research specimens provided written informed consent using an approved document.

## Supplemental Figure Legends

**Supplemental Figure 1. Immune profile of 37 GCPM samples by Bulk RNAseq; and expression of TIM3 ligand CEACAM1 and HMGB1 in GC tumors vs normal A.** Bulk RNA-seq analysis on notable immune markers on GCPM malignant ascites (n=37). Unsupervised hierarchical clustering was performed on the normalised RNA expression data of indicated genes. **B.** Increased TIM3 ligand CEACAM1 and HMGB1 in GC tumor tissues (n=408) compared to normal tissues (n=201) (http://gepia.cancer-pku.cn). **C.** Expression of CEACAM1 and HMGB1 was increased in GC tumor tissues compared to normal analyzed in GSE33335 cohort and GSE146996 cohort respectively.

**Supplemental Figure 2. Expression of TIM3 was increased in peritoneal metastases compared with primary tumors and adjacent normal from GC patients. A.** Expression of TIM3 was assessed by co-immunofluorescent staining of TIM3 and macrophage marker CD163 or TIM3 and tumor marker YAP1 in GC continuum. **B.** Association of TIM3 ligand Galectin-9, CEACAM1 and HMGB1 expression with GC patients’ survival in more than 600 advanced GC patients from the TCGA database (https://kmplot.com).

**Supplemental Figure 3. Expression of TIM3 is enriched in M2 macrophages and associated with notable M2 macrophage markers. A.** Expression of CD163 and CSF1R was analyzed in violin plot using scRNA-seq of unfractionated live cell mixture from 20 GCPM samples in all cell types. **B**. scRNA-seq analysis of unfractionated live cell mixture from 20 GCPM samples on notable immune checkpoint markers in all cell types in dot plots. **C.** Significant **c**orrelation between TIM3 and CD68 or PD-L1 was assessed from public GC dataset (http://gepia.cancer-pku.cn); D. Significant **c**orrelation between TIM3 and CD163 was revealed in three independent GC chort (GSE33335, GSE146996 and GSE237876). **E**. UMAP overview of single cell RNA-seq profiling cell type phenotypes (40 samples [>200,000 cells], comprising 3 peritoneal tumor samples [7,210 cells]) and TIM3 (*HAVCR2*) expression of across the cell types.

**Supplemental Figure 4. Bone marrow-derived monocyte/macrophages (mBM) repolarization and expression of TIM3, its ligand Galectin 9 and M2 macrophage markers in tumor cell induced M2 macrophages.** A, Schematic procedure to generate mBM M2 macrophages; B. Expression of TIM3, Galectin9 and M2 markers in mBM treated with murine tumor cell Kp-Luc2 condition medium (cGM) compared to normal growth medium (GM) by qRT-PCR.

**Supplemental Figure 5. Effects of TIM3 KO or TIM3 blockage in macrophages on T cell functions A.** CD3 and CD8T cell immune response (perforin production) was assessed by flow cytometry in mouse PBMCs co-culturing with mouse M2 macrophages RAW control and RAW two TIM3KO clones; **B**. CD45+ immune cells (containing T cells and M2 macrpophages) from malignant ascites of GA071924 GCPM patient were treated with or without anti-TIM3 neutralizing antibody for 2 days. The production of IFNg, GranB and Perforin from CD3 T cells were analyzed by flow cytometry; **C**, Target GA0518mCh-2 cells were seeded and co-cultured with different ratio of effector human PBMC in presence of co-cultured with U937 vector controls (U937mOvB) or U937 TIM3KO. Tumor cell viability was measured by luciferase activity assay. All experiments were performed in triplicate per sample.

**Supplemental Figure 6. Increased expression of CCL20 in GC tumor tissues; and rCCL20 stimulates phosphorylation of p90RSK1/2 in GA0518 cells. A.** Expression profile of identified cytokines from cytokine array (CCL4,CCL7,CCL20, IL-1A and IL-6) and their receptors is shown in dot plot of scRNAseq analysis with normalised expression levels of indicated genes from 20 GCPM specimen. **B**. Increased expression of CCL20 and its receptor CCR6 in GC tumor tissues (n=408) compared to normal tissues (n=201) (http://gepia.cancer-pku.cn). *p<0.05. **C**. Expression of CCL20, CCL1, CCL4 and CCL7 in tumor tissues compared with normal tissues was shown in GSE146996 cohort. **D**. Phosphorylation of p90RSK1/2, ERK1/2 and AKT three kinases was measured in GA0518 cells treated with U937 control or TIM3KO cell culture medium (cGM) or treated with ERK or p90RSK1/2 inhibitors GDC-0994 or BI-D1870 by Western blot; **E**. Phosphorylated and total p90RSK1/2 were detected in GA0518 cells treated with the U937 cGM in presence of GDC-0994 and rCCL20 by Western blot.

**Supplemental Figure 7. Effects of macrophage depletion and TIM3 blockade on tumor growth and metastases *in vivo***. **A-C.** Kp-Luc tumors were grown subcutaneously (sc) in C57BL/6 mice and treated with anti-TIM3 neutralizing antibody in macrophage depletion condition using Clodronate-liposome or Control-liposome; **D-F.** C57BL/6 mice bearing Kp-Luc2 tumors were treated with neutralizing antibodies aTIM3 or aPD-L1 or their combination; tumor growth was monitored weekly as described in Methods; **G&H.** In KP-Luc2 IP metastatic model, C57BL/6 mice were intraperitoneally injected with Kp-Luc2 tumor cells and then treated with aTIM3 and aPD1 neutralizing antibodies or combined with MMC as indicated in Figure 7F. Images of metastatic PM numbers per mouse was plotted(**G**); quantification of tumor burden/metastatic PM nodes was shown (**H**). P value embedded in the graph. **I.** CD8 T cell numbers was assessed by flowcytometry from treated six groups. P value is shown in the graph.

## References

1. Siegel RL, Kratzer TB, Wagle NS, Sung H, Jemal A. Cancer statistics, 2026. CA Cancer J Clin. 2026;76(1):e70043. doi: 10.3322/caac.70043. PubMed PMID: 41528114; PMCID: PMC12798275.

2. Gill RS, Al-Adra DP, Nagendran J, Campbell S, Shi X, Haase E, Schiller D. Treatment of gastric cancer with peritoneal carcinomatosis by cytoreductive surgery and HIPEC: a systematic review of survival, mortality, and morbidity. J Surg Oncol. 2011;104(6):692–8. Epub 20110628. doi: 10.1002/jso.22017. PubMed PMID: 21713780.

3. Sadeghi B, Arvieux C, Glehen O, Beaujard AC, Rivoire M, Baulieux J, Fontaumard E, Brachet A, Caillot JL, Faure JL, Porcheron J, Peix JL, Francois Y, Vignal J, Gilly FN. Peritoneal carcinomatosis from non-gynecologic malignancies: results of the EVOCAPE 1 multicentric prospective study. Cancer. 2000;88(2):358–63. doi: 10.1002/(sici)1097-0142(20000115)88:2<358::aid-cncr16>3.0.co;2-o. PubMed PMID: 10640968.

4. Pyrhönen S, Kuitunen T, Nyandoto P, Kouri M. Randomised comparison of fluorouracil, epidoxorubicin and methotrexate (FEMTX) plus supportive care with supportive care alone in patients with non-resectable gastric cancer. British journal of cancer. 1995;71(3):587–91. PubMed PMID: 7533517; PMCID: PMC2033628.

5. Kobayashi D, Kodera Y. Intraperitoneal chemotherapy for gastric cancer with peritoneal metastasis. Gastric cancer : official journal of the International Gastric Cancer Association and the Japanese Gastric Cancer Association. 2016. doi: 10.1007/s10120-016-0662-9. PubMed PMID: 27803990.

6. Satoi S, Fujii T, Yanagimoto H, Motoi F, Kurata M, Takahara N, Yamada S, Yamamoto T, Mizuma M, Honda G, Isayama H, Unno M, Kodera Y, Ishigami H, Kon M. Multicenter Phase II Study of Intravenous and Intraperitoneal Paclitaxel With S-1 for Pancreatic Ductal Adenocarcinoma Patients With Peritoneal Metastasis. Annals of surgery. 2016. doi: 10.1097/SLA.0000000000001705. PubMed PMID: 26982692.

7. Shi C, Yang B, Chen Q, Yang J, Fan N. Retrospective analysis of adjuvant intraperitoneal chemotherapy effect prognosis of resectable gastric cancer. Oncology. 2011;80(5-6):289–95. Epub 20110718. doi: 10.1159/000329075. PubMed PMID: 21778768.

8. Sauer N, Janicka N, Szlasa W, Skinderowicz B, Kolodzinska K, Dwernicka W, Oslizlo M, Kulbacka J, Novickij V, Karlowicz-Bodalska K. TIM-3 as a promising target for cancer immunotherapy in a wide range of tumors. Cancer Immunol Immunother. 2023;72(11):3405–25. Epub 20230811. doi: 10.1007/s00262-023-03516-1. PubMed PMID: 37567938; PMCID: PMC10576709.

9. Monney L, Sabatos CA, Gaglia JL, Ryu A, Waldner H, Chernova T, Manning S, Greenfield EA, Coyle AJ, Sobel RA, Freeman GJ, Kuchroo VK. Th1-specific cell surface protein Tim-3 regulates macrophage activation and severity of an autoimmune disease. Nature. 2002;415(6871):536–41. doi: 10.1038/415536a. PubMed PMID: 11823861.

10. Gao X, Zhu Y, Li G, Huang H, Zhang G, Wang F, Sun J, Yang Q, Zhang X, Lu B. TIM-3 expression characterizes regulatory T cells in tumor tissues and is associated with lung cancer progression. PLoS One. 2012;7(2):e30676. Epub 20120217. doi: 10.1371/journal.pone.0030676. PubMed PMID: 22363469; PMCID: PMC3281852.

11. Severson JJ, Serracino HS, Mateescu V, Raeburn CD, McIntyre RC, Jr., Sams SB, Haugen BR, French JD. PD-1+Tim-3+ CD8+ T Lymphocytes Display Varied Degrees of Functional Exhaustion in Patients with Regionally Metastatic Differentiated Thyroid Cancer. Cancer Immunol Res. 2015;3(6):620–30. Epub 20150219. doi: 10.1158/2326-6066.CIR-14-0201. PubMed PMID: 25701326; PMCID: PMC4457654.

12. Acharya N, Sabatos-Peyton C, Anderson AC. Tim-3 finds its place in the cancer immunotherapy landscape. J Immunother Cancer. 2020;8(1). doi: 10.1136/jitc-2020-000911. PubMed PMID: 32601081; PMCID: PMC7326247.

13. Zhang C, Xu L, Ma Y, Huang Y, Zhou L, Le H, Chen Z. Increased TIM-3 expression in tumor-associated macrophages predicts a poorer prognosis in non-small cell lung cancer: a retrospective cohort study. J Thorac Dis. 2023;15(3):1433–44. Epub 20230327. doi: 10.21037/jtd-23-227. PubMed PMID: 37065598; PMCID: PMC10089863.

14. Vanmeerbeek I, Naulaerts S, Sprooten J, Laureano RS, Govaerts J, Trotta R, Pretto S, Zhao S, Cafarello ST, Verelst J, Jacquemyn M, Pociupany M, Boon L, Schlenner SM, Tejpar S, Daelemans D, Mazzone M, Garg AD. Targeting conserved TIM3(+)VISTA(+) tumor-associated macrophages overcomes resistance to cancer immunotherapy. Sci Adv. 2024;10(29):eadm8660. Epub 20240719. doi: 10.1126/sciadv.adm8660. PubMed PMID: 39028818; PMCID: PMC11259173.

15. Dixon KO, Tabaka M, Schramm MA, Xiao S, Tang R, Dionne D, Anderson AC, Rozenblatt-Rosen O, Regev A, Kuchroo VK. TIM-3 restrains anti-tumour immunity by regulating inflammasome activation. Nature. 2021;595(7865):101–6. Epub 20210609. doi: 10.1038/s41586-021-03626-9. PubMed PMID: 34108686; PMCID: PMC8627694.

16. Wang Z, Yin N, Zhang Z, Zhang Y, Zhang G, Chen W. Upregulation of T-cell Immunoglobulin and Mucin-Domain Containing-3 (Tim-3) in Monocytes/Macrophages Associates with Gastric Cancer Progression. Immunol Invest. 2017;46(2):134–48. Epub 20161202. doi: 10.1080/08820139.2016.1229790. PubMed PMID: 27911104.

17. Ausejo-Mauleon I, Labiano S, de la Nava D, Laspidea V, Zalacain M, Marrodan L, Garcia-Moure M, Gonzalez-Huarriz M, Hervas-Corpion I, Dhandapani L, Vicent S, Collantes M, Penuelas I, Becher OJ, Filbin MG, Jiang L, Labelle J, de Biagi-Junior CAO, Nazarian J, Laternser S, Phoenix TN, van der Lugt J, Kranendonk M, Hoogendijk R, Mueller S, De Andrea C, Anderson AC, Guruceaga E, Koschmann C, Yadav VN, Gallego Perez-Larraya J, Patino-Garcia A, Pastor F, Alonso MM. TIM-3 blockade in diffuse intrinsic pontine glioma models promotes tumor regression and antitumor immune memory. Cancer Cell. 2023;41(11):1911–26 e8. Epub 20231005. doi: 10.1016/j.ccell.2023.09.001. PubMed PMID: 37802053; PMCID: PMC10644900.

18. Wang R, Song S, Harada K, Ghazanfari Amlashi F, Badgwell B, Pizzi MP, Xu Y, Zhao W, Dong X, Jin J, Wang Y, Scott A, Ma L, Huo L, Vicente D, Blum Murphy M, Shanbhag N, Tatlonghari G, Thomas I, Rogers J, Kobayashi M, Vykoukal J, Estrella JS, Roy-Chowdhuri S, Han G, Zhang S, Mao X, Song X, Zhang J, Gu J, Johnson RL, Calin GA, Peng G, Lee JS, Hanash SM, Futreal A, Wang Z, Wang L, Ajani JA. Multiplex profiling of peritoneal metastases from gastric adenocarcinoma identified novel targets and molecular subtypes that predict treatment response. Gut. 2020;69(1):18–31. Epub 20190606. doi: 10.1136/gutjnl-2018-318070. PubMed PMID: 31171626; PMCID: PMC6943252.

19. Zhao JJ, Ong CJ, Srivastava S, Chia DKA, Ma H, Huang K, Sheng T, Ramnarayanan K, Ong X, Tay ST, Hagihara T, Tan ALK, Teo MCC, Tan QX, Ng G, Tan JW, Ng MCH, Gwee YX, Walsh R, Law JH, Shabbir A, Kim G, Tay Y, Her Z, Leoncini G, Teh BT, Hong JH, Tay RYK, Teo CB, Dings MPG, Bijlsma M, Lum JHY, Mathur S, Pietrantonio F, Blum SM, van Laarhoven H, Klempner SJ, Yong WP, So JBY, Chen Q, Tan P, Sundar R. Spatially Resolved Niche and Tumor Microenvironmental Alterations in Gastric Cancer Peritoneal Metastases. Gastroenterology. 2024;167(7):1384–98 e4. Epub 20240813. doi: 10.1053/j.gastro.2024.08.007. PubMed PMID: 39147169.

20. Fan Y, Li Y, Yao X, Jin J, Scott A, Liu B, Wang S, Huo L, Wang Y, Wang R, Pool Pizzi M, Ma L, Shao S, Sewastjanow-Silva M, Waters R, Chatterjee D, Liu B, Shanbhag N, Peng G, Calin GA, Mazur PK, Hanash SM, Ishizawa J, Hirata Y, Nagano O, Wang Z, Wang L, Xian W, McKeon F, Ajani JA, Song S. Epithelial SOX9 drives progression and metastases of gastric adenocarcinoma by promoting immunosuppressive tumour microenvironment. Gut. 2023;72(4):624–37. Epub 20220824. doi: 10.1136/gutjnl-2021-326581. PubMed PMID: 36002248.

21. Kumar V, Ramnarayanan K, Sundar R, Padmanabhan N, Srivastava S, Koiwa M, Yasuda T, Koh V, Huang KK, Tay ST, Ho SWT, Tan ALK, Ishimoto T, Kim G, Shabbir A, Chen Q, Zhang B, Xu S, Lam KP, Lum HYJ, Teh M, Yong WP, So JBY, Tan P. Single-Cell Atlas of Lineage States, Tumor Microenvironment, and Subtype-Specific Expression Programs in Gastric Cancer. Cancer Discov. 2022;12(3):670–91. doi: 10.1158/2159-8290.CD-21-0683. PubMed PMID: 34642171; PMCID: PMC9394383.

22. Song S, Xu Y, Huo L, Zhao S, Wang R, Li Y, Scott AW, Pizzi MP, Wang Y, Fan Y, Harada K, Jin J, Ma L, Yao X, Shanbhag ND, Gan Q, Roy-Chowdhuri S, Badgwell BD, Wang Z, Wang L, Ajani JA. Patient-derived cell lines and orthotopic mouse model of peritoneal carcinomatosis recapitulate molecular and phenotypic features of human gastric adenocarcinoma. J Exp Clin Cancer Res. 2021;40(1):207. Epub 20210623. doi: 10.1186/s13046-021-02003-8. PubMed PMID: 34162421; PMCID: PMC8223395.

23. Chow A, Schad S, Green MD, Hellmann MD, Allaj V, Ceglia N, Zago G, Shah NS, Sharma SK, Mattar M, Chan J, Rizvi H, Zhong H, Liu C, Bykov Y, Zamarin D, Shi H, Budhu S, Wohlhieter C, Uddin F, Gupta A, Khodos I, Waninger JJ, Qin A, Markowitz GJ, Mittal V, Balachandran V, Durham JN, Le DT, Zou W, Shah SP, McPherson A, Panageas K, Lewis JS, Perry JSA, de Stanchina E, Sen T, Poirier JT, Wolchok JD, Rudin CM, Merghoub T. Tim-4(+) cavity-resident macrophages impair anti-tumor CD8(+) T cell immunity. Cancer Cell. 2021;39(7):973–88 e9. Epub 20210610. doi: 10.1016/j.ccell.2021.05.006. PubMed PMID: 34115989; PMCID: PMC9115604.

24. Kadomoto S, Izumi K, Mizokami A. The CCL20-CCR6 Axis in Cancer Progression. Int J Mol Sci. 2020;21(15). Epub 20200722. doi: 10.3390/ijms21155186. PubMed PMID: 32707869; PMCID: PMC7432448.

25. Ranasinghe R, Eri R. Modulation of the CCR6-CCL20 Axis: A Potential Therapeutic Target in Inflammation and Cancer. Medicina (Kaunas). 2018;54(5). Epub 20181116. doi: 10.3390/medicina54050088. PubMed PMID: 30453514; PMCID: PMC6262638.

26. Papadopoulos KP, Gluck L, Martin LP, Olszanski AJ, Tolcher AW, Ngarmchamnanrith G, Rasmussen E, Amore BM, Nagorsen D, Hill JS, Stephenson J, Jr. First-in-Human Study of AMG 820, a Monoclonal Anti-Colony-Stimulating Factor 1 Receptor Antibody, in Patients with Advanced Solid Tumors. Clin Cancer Res. 2017;23(19):5703–10. Epub 20170627. doi: 10.1158/1078-0432.CCR-16-3261. PubMed PMID: 28655795.

27. Okazaki S, Shintani S, Hirata Y, Suina K, Semba T, Yamasaki J, Umene K, Ishikawa M, Saya H, Nagano O. Synthetic lethality of the ALDH3A1 inhibitor dyclonine and xCT inhibitors in glutathione deficiency-resistant cancer cells. Oncotarget. 2018;9(73):33832–43. Epub 20180918. doi: 10.18632/oncotarget.26112. PubMed PMID: 30333913; PMCID: PMC6173468.

28. Tanaka Y, Chiwaki F, Kojima S, Kawazu M, Komatsu M, Ueno T, Inoue S, Sekine S, Matsusaki K, Matsushita H, Boku N, Kanai Y, Yatabe Y, Sasaki H, Mano H. Multi-omic profiling of peritoneal metastases in gastric cancer identifies molecular subtypes and therapeutic vulnerabilities. Nat Cancer. 2021;2(9):962–77. Epub 20210816. doi: 10.1038/s43018-021-00240-6. PubMed PMID: 35121863.

29. Verwaal VJ, van Ruth S, de Bree E, van Sloothen GW, van Tinteren H, Boot H, Zoetmulder FA. Randomized trial of cytoreduction and hyperthermic intraperitoneal chemotherapy versus systemic chemotherapy and palliative surgery in patients with peritoneal carcinomatosis of colorectal cancer. J Clin Oncol. 2003;21(20):3737–43. doi: 10.1200/JCO.2003.04.187. PubMed PMID: 14551293.

30. van Driel WJ, Koole SN, Sonke GS. Hyperthermic Intraperitoneal Chemotherapy in Ovarian Cancer. N Engl J Med. 2018;378(14):1363–4. doi: 10.1056/NEJMc1802033. PubMed PMID: 29617590.

31. Bayik D, Lathia JD. Cancer stem cell-immune cell crosstalk in tumour progression. Nat Rev Cancer. 2021;21(8):526–36. Epub 20210608. doi: 10.1038/s41568-021-00366-w. PubMed PMID: 34103704; PMCID: PMC8740903.

32. Bied M, Ho WW, Ginhoux F, Bleriot C. Roles of macrophages in tumor development: a spatiotemporal perspective. Cell Mol Immunol. 2023;20(9):983–92. Epub 20230710. doi: 10.1038/s41423-023-01061-6. PubMed PMID: 37429944; PMCID: PMC10468537.

33. Song H, Wang T, Tian L, Bai S, Chen L, Zuo Y, Xue Y. Macrophages on the Peritoneum are involved in Gastric Cancer Peritoneal Metastasis. J Cancer. 2019;10(22):5377–87. Epub 20190829. doi: 10.7150/jca.31787. PubMed PMID: 31632482; PMCID: PMC6775704.

34. Sathe A, Mason K, Grimes SM, Zhou Z, Lau BT, Bai X, Su A, Tan X, Lee H, Suarez CJ, Nguyen Q, Poultsides G, Zhang NR, Ji HP. Colorectal Cancer Metastases in the Liver Establish Immunosuppressive Spatial Networking between Tumor-Associated SPP1+ Macrophages and Fibroblasts. Clin Cancer Res. 2023;29(1):244–60. doi: 10.1158/1078-0432.CCR-22-2041. PubMed PMID: 36239989; PMCID: PMC9811165.

35. Wang R, Song S, Qin J, Yoshimura K, Peng F, Chu Y, Li Y, Fan Y, Jin J, Dang M, Dai E, Pei G, Han G, Hao D, Li Y, Chatterjee D, Harada K, Pizzi MP, Scott AW, Tatlonghari G, Yan X, Xu Z, Hu C, Mo S, Shanbhag N, Lu Y, Sewastjanow-Silva M, Fouad Abdelhakeem AA, Peng G, Hanash SM, Calin GA, Yee C, Mazur P, Marsden AN, Futreal A, Wang Z, Cheng X, Ajani JA, Wang L. Evolution of immune and stromal cell states and ecotypes during gastric adenocarcinoma progression. Cancer Cell. 2023;41(8):1407–26 e9. Epub 20230706. doi: 10.1016/j.ccell.2023.06.005. PubMed PMID: 37419119; PMCID: PMC10528152.

36. Samaniego R, Gutierrez-Gonzalez A, Gutierrez-Seijo A, Sanchez-Gregorio S, Garcia-Gimenez J, Mercader E, Marquez-Rodas I, Aviles JA, Relloso M, Sanchez-Mateos P. CCL20 Expression by Tumor-Associated Macrophages Predicts Progression of Human Primary Cutaneous Melanoma. Cancer Immunol Res. 2018;6(3):267–75. Epub 20180123. doi: 10.1158/2326-6066.CIR-17-0198. PubMed PMID: 29362221.

37. Romeo Y, Zhang X, Roux PP. Regulation and function of the RSK family of protein kinases. Biochem J. 2012;441(2):553–69. doi: 10.1042/BJ20110289. PubMed PMID: 22187936.

38. Popovici RM, Krause MS, Germeyer A, Strowitzki T, von Wolff M. Galectin-9: a new endometrial epithelial marker for the mid- and late-secretory and decidual phases in humans. J Clin Endocrinol Metab. 2005;90(11):6170–6. Epub 20050816. doi: 10.1210/jc.2004-2529. PubMed PMID: 16105962.

39. Fan Y, Song S, Li Y, Dhar SS, Jin J, Yoshimura K, Yao X, Wang R, Scott AW, Pizzi MP, Wu J, Ma L, Calin GA, Hanash S, Wang L, Curran M, Ajani JA. Galectin-3 Cooperates with CD47 to Suppress Phagocytosis and T-cell Immunity in Gastric Cancer Peritoneal Metastases. Cancer Res. 2023;83(22):3726–38. doi: 10.1158/0008-5472.CAN-23-0783. PubMed PMID: 37738407; PMCID: PMC10843008.

40. Ajani JA, Xu Y, Huo L, Wang R, Li Y, Wang Y, Pizzi MP, Scott A, Harada K, Ma L, Yao X, Jin J, Zhao W, Dong X, Badgwell BD, Shanbhag N, Tatlonghari G, Estrella JS, Roy-Chowdhuri S, Kobayashi M, Vykoukal JV, Hanash SM, Calin GA, Peng G, Lee JS, Johnson RL, Wang Z, Wang L, Song S. YAP1 mediates gastric adenocarcinoma peritoneal metastases that are attenuated by YAP1 inhibition. Gut. 2021;70(1):55–66. Epub 20200427. doi: 10.1136/gutjnl-2019-319748. PubMed PMID: 32345613; PMCID: PMC9832914.

41. Wang R, Dang M, Harada K, Han G, Wang F, Pool Pizzi M, Zhao M, Tatlonghari G, Zhang S, Hao D, Lu Y, Zhao S, Badgwell BD, Blum Murphy M, Shanbhag N, Estrella JS, Roy-Chowdhuri S, Abdelhakeem AAF, Wang Y, Peng G, Hanash S, Calin GA, Song X, Chu Y, Zhang J, Li M, Chen K, Lazar AJ, Futreal A, Song S, Ajani JA, Wang L. Single-cell dissection of intratumoral heterogeneity and lineage diversity in metastatic gastric adenocarcinoma. Nat Med. 2021;27(1):141–51. Epub 20210104. doi: 10.1038/s41591-020-1125-8. PubMed PMID: 33398161; PMCID: PMC8074162.

