## Supplemental Figures 1-7 for "TIM3^+^ Tumor Associated M2 Macrophages Impair Antitumor T Cell Immunity and Promote Gastric Cancer Progression and Peritoneal Metastasis"

### Supplemental Figure 1.

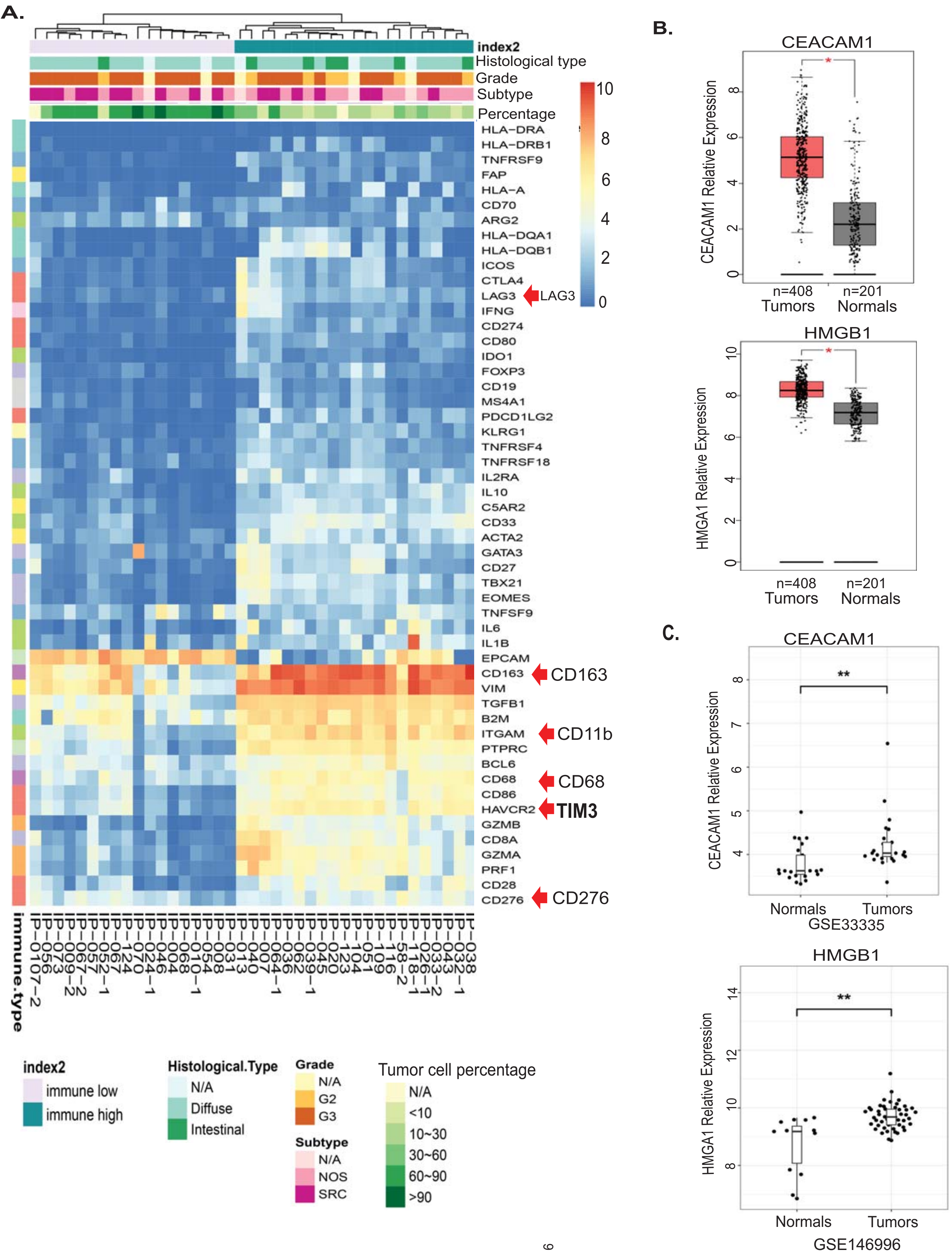

#### Supplemental Figure 2

A.

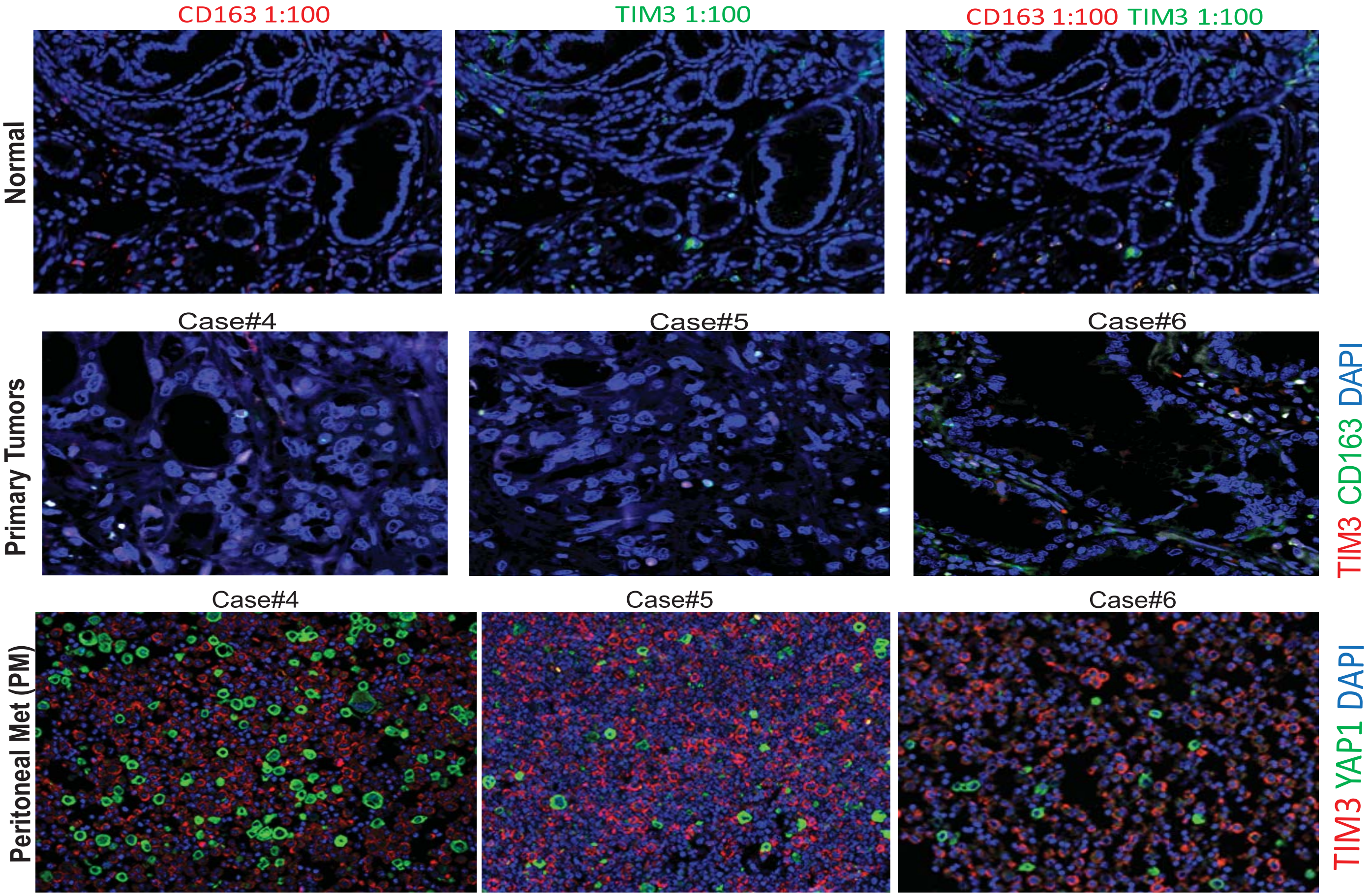


B.

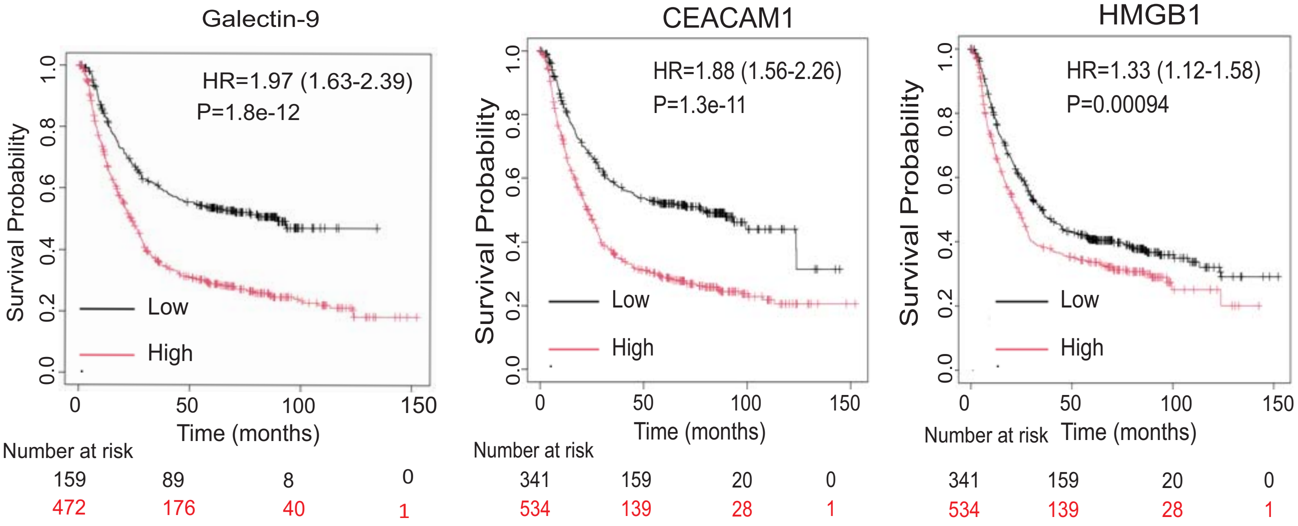

**Supplmental Figure 3.**

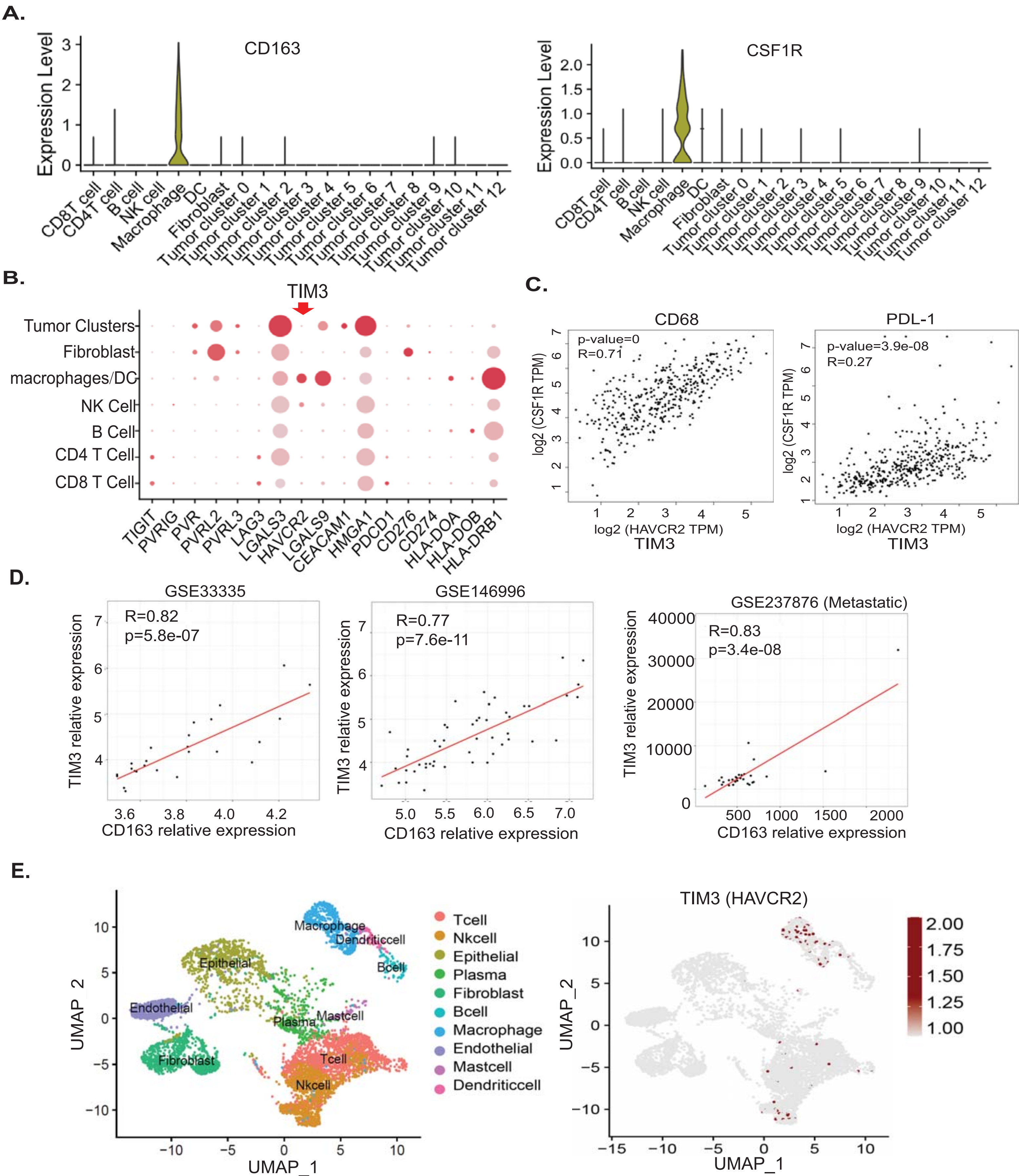


**Peritoneal tumor samples only**

n=3 PM samples; 7,210 cells

#### Supplemental Figure 4

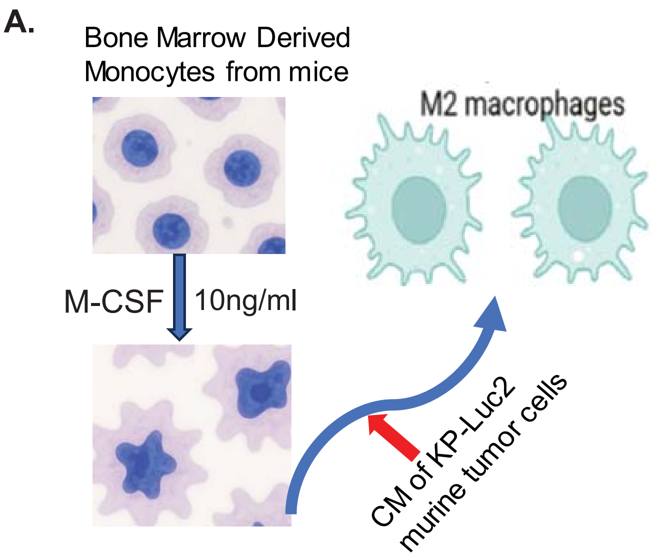


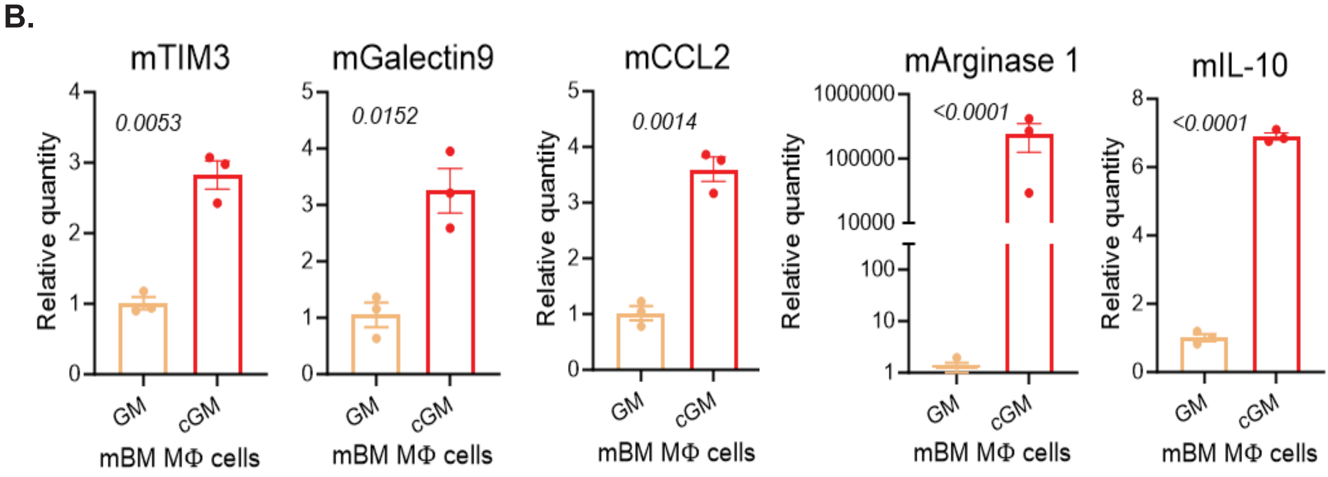

Supplemental Figure 5

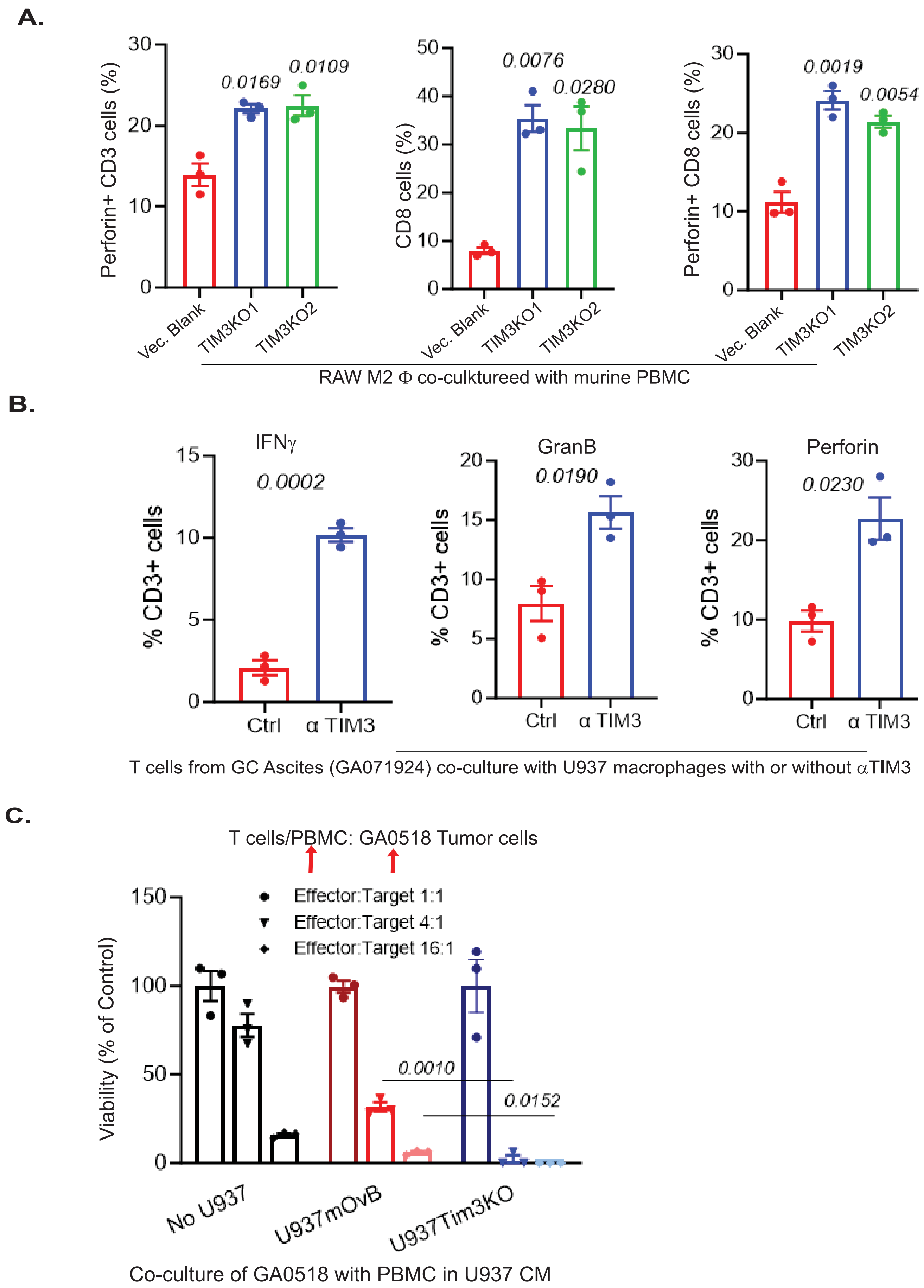

**Supplemental Figure 6.**

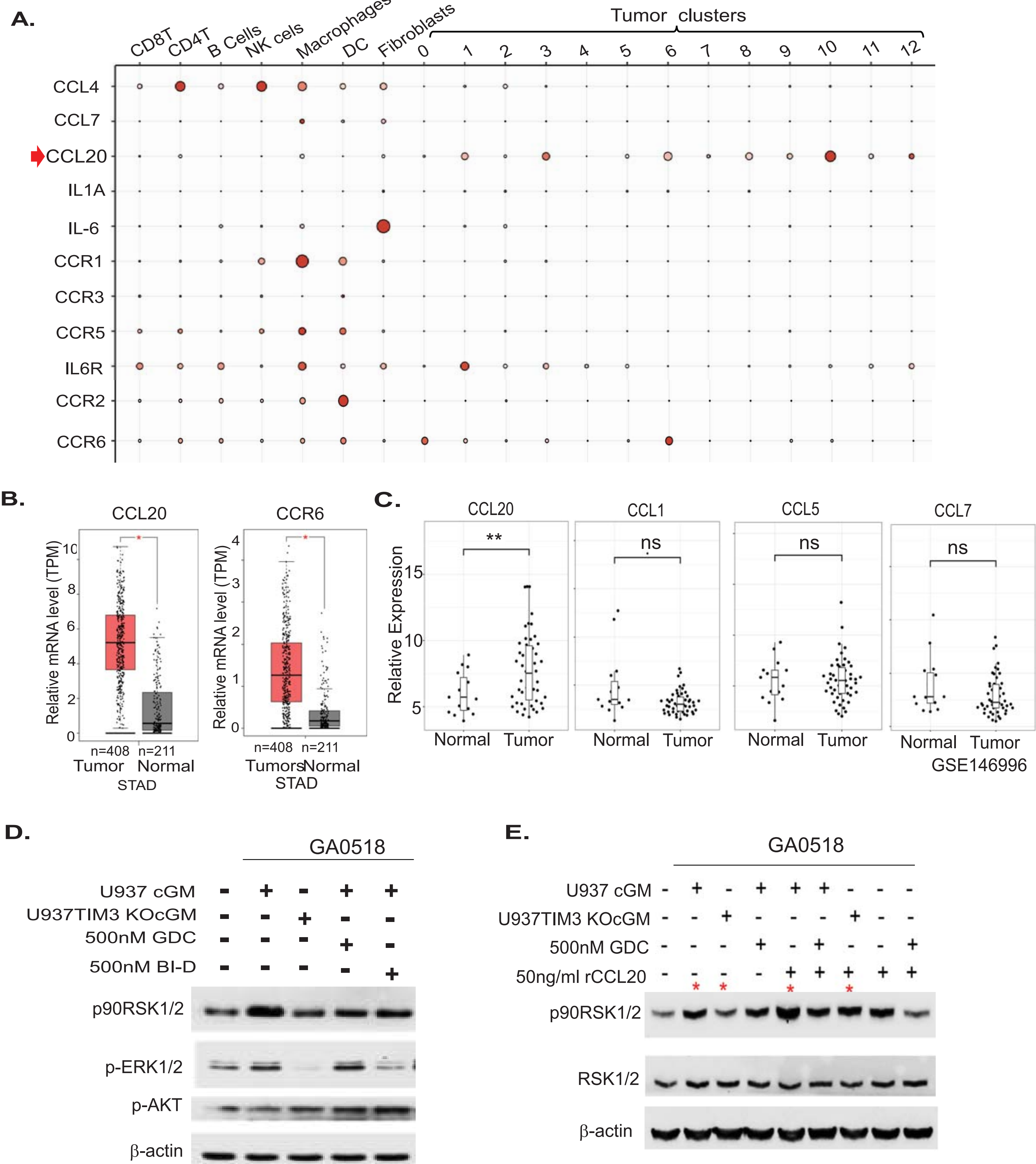

**Supplemental Figure 7.**

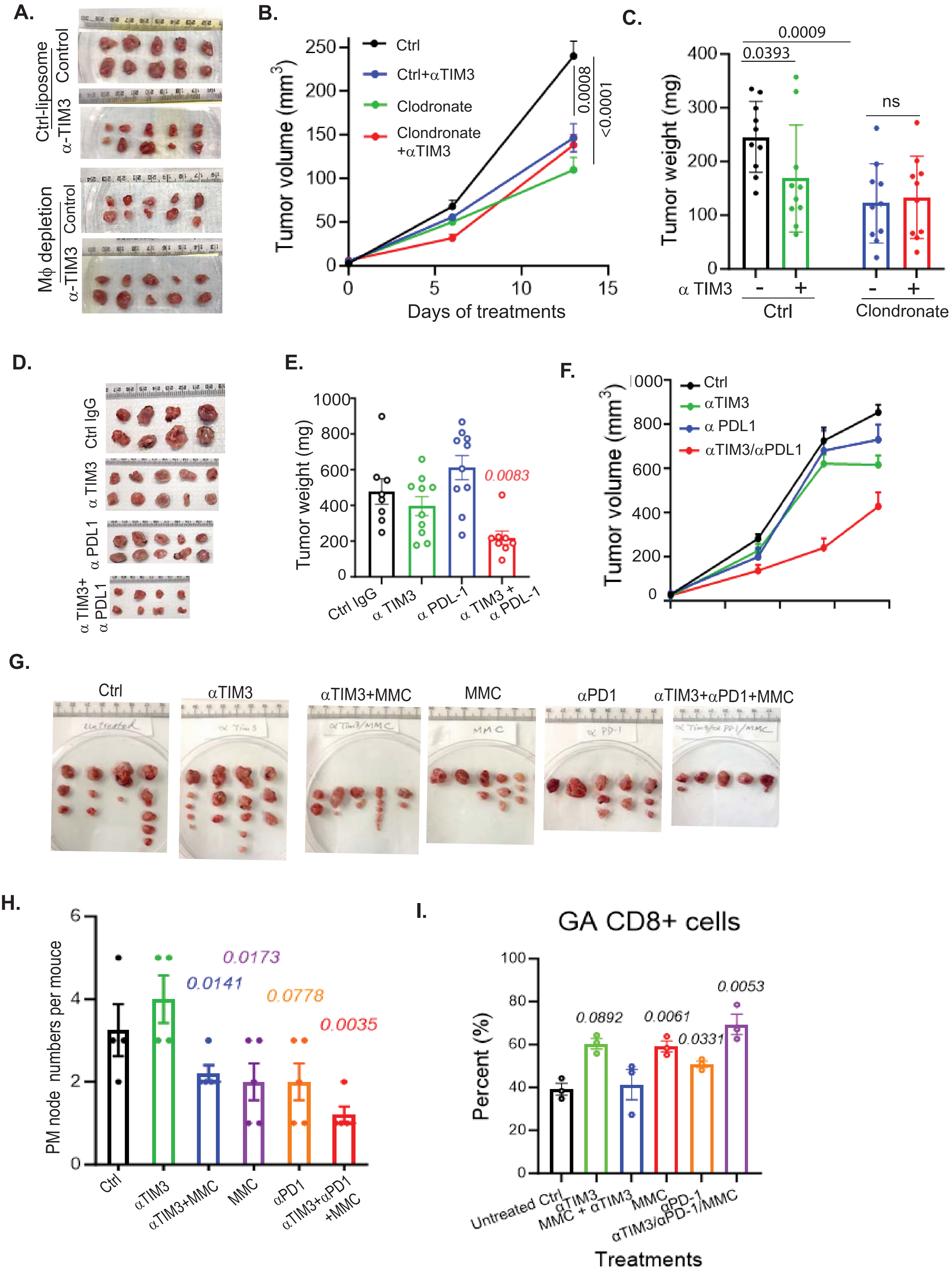
