## Supplemental Table 1 for "TIM3^+^ Tumor Associated M2 Macrophages Impair Antitumor T Cell Immunity and Promote Gastric Cancer Progression and Peritoneal Metastasis"

**Supplemental Table 1. The cell subtypes and TIM-3 and its relative proteins in gastric ascites of GAC patients**

| Specimen # | Subtypes | | | | | | TIM-3 | | | | | |
|---|---|---|---|---|---|---|---|---|---|---|---|---|
| | EpCAM | CD45 | CD3 | CD4 | CD8 | CD163 | EpCAM | CD45 | CD3 | CD4 | CD8 | CD163 |
| 1 | 1.58 | 51.90 | 40.90 | 29.70 | 26.40 | 26.60 | 13.30 | 55.90 | 0.12 | 0.46 | 0.41 | 94.40 |
| 2 | 0.64 | 65.10 | 45.20 | 3.08 | 33.30 | 1.86 | 0.94 | 3.89 | 0.05 | 0.94 | 0.17 | 77.00 |
| 3 | 52.30 | 30.50 | 14.30 | 12.00 | 19.10 | 4.62 | 69.40 | 19.90 | 0.13 | 0.90 | 0.11 | 67.20 |
| 4 | 91.10 | 1.94 | 23.10 | 17.70 | 10.70 | 6.21 | 0.17 | 10.10 | 0.03 | 0.34 | 0.28 | 83.90 |
| 5 | 42.90 | 12.60 | 24.60 | 20.20 | 11.20 | 19.40 | 74.20 | 57.70 | 0.29 | 1.84 | 0.37 | 91.90 |
| 6 | 0.16 | 90.40 | 45.80 | 44.20 | 18.10 | 10.10 | 5.98 | 25.40 | 0.06 | 0.31 | 0.13 | 96.60 |
| 7 | 6.76 | 47.70 | 42.10 | 69.80 | 6.27 | 26.00 | 9.23 | 57.30 | 0.81 | 1.51 | 4.70 | 99.50 |
| 8 | 2.39 | 76.80 | 68.40 | 64.00 | 11.80 | 4.74 | 26.80 | 17.10 | 0.21 | 0.23 | 0.78 | 97.20 |
| 9 | 1.10 | 82.80 | 28.00 | 35.70 | 41.30 | 26.40 | 16.20 | 62.00 | 1.37 | 5.48 | 2.31 | 98.60 |
| 10 | 84.00 | 1.89 | 43.40 | 2.89 | 1.43 | 59.90 | 24.20 | 84.90 | 0.17 | 17.40 | 2.72 | 98.10 |
| 11 | 0.27 | 98.40 | 41.90 | 35.40 | 28.90 | 47.00 | 51.00 | 85.00 | 1.06 | 2.36 | 2.28 | 99.80 |
| 12 | 4.52 | 90.80 | 76.30 | 61.40 | 15.10 | 0.66 | 2.55 | 1.67 | 0.12 | 0.10 | 0.22 | 84.60 |
| 13 | 29.40 | 12.20 | 44.30 | 12.10 | 16.50 | 7.41 | 34.50 | 20.40 | 0.84 | 3.08 | 2.82 | 67.50 |
| 14 | 6.17 | 79.00 | 60.10 | 58.40 | 16.40 | 3.49 | 0.34 | 11.70 | 0.14 | 0.16 | 0.13 | 81.60 |
| 15 | 0.04 | 34.40 | 69.80 | 42.70 | 15.90 | 6.37 | 7.21 | 15.00 | 0.31 | 0.28 | 0.75 | 86.80 |
| 16 | 0.06 | 87.90 | 22.50 | 26.70 | 24.60 | 22.70 | 12.50 | 58.70 | 1.03 | 1.36 | 2.62 | 95.00 |
| 17 | 0.01 | 15.80 | 41.90 | 31.70 | 8.56 | 1.80 | 0.00 | 1.93 | 0.07 | 0.06 | 0.21 | 43.40 |
| 18 | 39.90 | 7.86 | 7.62 | 12.30 | 3.82 | 17.90 | 0.34 | 63.20 | 11.30 | 5.11 | 10.90 | 99.00 |
| 19 | 1.31 | 8.97 | 12.30 | 28.30 | 4.59 | 8.27 | 0.51 | 22.40 | 3.02 | 2.27 | 2.54 | 97.60 |
| 20 | 85.00 | 1.10 | 0.67 | 0.66 | 1.04 | 61.60 | 0.12 | 90.00 | 0.00 | 1.94 | 1.29 | 99.00 |
| 21 | 0.61 | 20.30 | 33.90 | 2.69 | 17.40 | 4.91 | 0.77 | 17.10 | 1.11 | 13.80 | 0.55 | 82.90 |
| 22 | 3.77 | 8.37 | 39.50 | 3.53 | 18.20 | 5.35 | 7.94 | 14.70 | 4.06 | 3.24 | 10.50 | 51.90 |
| 23 | 0.01 | 13.60 | 80.10 | 5.25 | 27.00 | 31.70 | 22.70 | 77.60 | 2.88 | 17.50 | 9.36 | 99.20 |
| 24 | 0.03 | 16.30 | 59.80 | 9.42 | 29.60 | 6.06 | 0.00 | 21.90 | 0.69 | 1.86 | 1.21 | 83.90 |
| 25 | 0.50 | 6.26 | 39.40 | 5.96 | 7.67 | 2.39 | 0.00 | 6.11 | 0.06 | 1.34 | 0.10 | 64.30 |
| 26 | 5.01 | 10.30 | 37.10 | 11.80 | 10.40 | 31.60 | 0.37 | 80.30 | 1.45 | 8.84 | 0.18 | 99.80 |
| 27 | 21.90 | 9.21 | 51.70 | 8.89 | 8.76 | 2.06 | 0.06 | 5.30 | 0.35 | 2.05 | 0.00 | 59.30 |

| | | | | | | | | | | | | |
|---|---|---|---|---|---|---|---|---|---|---|---|---|
| 28 | 0.08 | 11.50 | 47.20 | 8.31 | 19.10 | 4.77 | 0.00 | 25.70 | 1.17 | 6.05 | 0.27 | 82.30 |
| 29 | 29.50 | 13.70 | 22.20 | 16.30 | 12.40 | 7.61 | 0.62 | 37.60 | 0.64 | 5.87 | 0.28 | 73.70 |
| 30 | 1.51 | 1.86 | 69.90 | 2.80 | 24.10 | 42.10 | 3.64 | 48.80 | 3.02 | 6.70 | 10.90 | 97.60 |
| 31 | 1.97 | 23.80 | 26.10 | 19.90 | 10.10 | 23.90 | 0.22 | 58.60 | 3.35 | 9.15 | 2.96 | 98.20 |
| 32 | 0.01 | 12.50 | 58.70 | 3.14 | 17.10 | 8.17 | 9.09 | 0.31 | 2.78 | 7.04 | 10.10 | 87.80 |
| 33 | 2.43 | 14.50 | 49.40 | 4.33 | 10.90 | 9.29 | 0.56 | 12.00 | 2.04 | 3.86 | 5.83 | 88.20 |
| 34 | 1.18 | 7.44 | 69.20 | 11.80 | 38.70 | 15.70 | 0.14 | 24.80 | 1.94 | 0.63 | 7.69 | 95.90 |
| 35 | 43.60 | 2.08 | 73.10 | 4.58 | 26.10 | 21.50 | 0.27 | 81.80 | 4.97 | 1.40 | 5.13 | 99.60 |
| 36 | 13.00 | 13.30 | 57.10 | 8.61 | 12.60 | 9.13 | 1.32 | 40.50 | 7.01 | 4.28 | 11.50 | 94.70 |
| 37 | 59.80 | 3.14 | 43.30 | 9.02 | 15.60 | 36.20 | 11.50 | 63.60 | 2.80 | 35.40 | 2.67 | 96.30 |
| 38 | 16.30 | 6.28 | 75.40 | 8.08 | 9.54 | 5.80 | 0.36 | 32.60 | 0.93 | 25.70 | 3.27 | 98.90 |
| 39 | 3.06 | 22.60 | 43.30 | 4.29 | 21.60 | 0.83 | 0.25 | 4.08 | 0.16 | 0.58 | 0.69 | 51.00 |
| 40 | 0.35 | 13.10 | 75.20 | 5.57 | 29.80 | 21.30 | 0.10 | 26.80 | 2.67 | 26.70 | 2.63 | 99.90 |
| 41 | 0.00 | 1.71 | 70.70 | 5.21 | 36.30 | 22.60 | 7.14 | 35.50 | 1.68 | 5.15 | 1.47 | 98.00 |
| 42 | 1.70 | 1.49 | 39.20 | 5.44 | 38.40 | 36.30 | 4.04 | 79.50 | 1.80 | 2.63 | 0.96 | 99.20 |
| 43 | 3.06 | 22.60 | 43.30 | 4.29 | 21.60 | 0.83 | 0.25 | 4.08 | 0.16 | 0.58 | 0.59 | 51.00 |
| 44 | 3.53 | 2.95 | 71.20 | 9.60 | 30.30 | 2.81 | 0.56 | 5.39 | 0.06 | 0.27 | 0.05 | 93.20 |
| 45 | 6.29 | 6.48 | 69.50 | 8.82 | 28.80 | 4.17 | 0.27 | 8.88 | 0.13 | 0.89 | 0.03 | 45.70 |
| 46 | 0.47 | 12.10 | 51.60 | 8.74 | 6.38 | 3.78 | 1.54 | 5.00 | 0.10 | 0.10 | 0.84 | 91.70 |
| 47 | 0.28 | 25.90 | 25.40 | 3.62 | 12.50 | 2.71 | 0.00 | 6.62 | 1.80 | 11.10 | 1.21 | 88.90 |
| 48 | 0.03 | 3.71 | 57.10 | 10.10 | 23.40 | 0.10 | 0.00 | 0.77 | 0.06 | 0.28 | 0.15 | 41.70 |
| 49 | 0.03 | 1.50 | 48.90 | 5.33 | 10.10 | 2.70 | 17.50 | 13.30 | 1.26 | 4.26 | 0.54 | 82.40 |
| 50 | 0.61 | 4.50 | 56.40 | 18.70 | 23.20 | 9.22 | 0.75 | 11.00 | 0.37 | 9.49 | 0.19 | 83.60 |
| 51 | 56.60 | 3.70 | 3.81 | 0.74 | 0.73 | 5.07 | 2.47 | 72.10 | 0.87 | 13.70 | 0.85 | 95.90 |
| 52 | 15.40 | 16.30 | 13.50 | 5.79 | 11.80 | 56.70 | 49.70 | 69.90 | 2.57 | 27.10 | 0.71 | 98.40 |
| 53 | 0.06 | 4.36 | 34.10 | 3.15 | 11.90 | 2.49 | 0.00 | 6.10 | 0.32 | 2.64 | 0.37 | 93.60 |
| 54 | 9.95 | 12.10 | 62.70 | 3.44 | 16.50 | 2.43 | 0.34 | 7.77 | 0.62 | 14.60 | 1.06 | 91.00 |
| **Specimen #** | | | **PD-1** | | | | | | **PD-L1** | | | |
| | EpCAM | CD45 | CD3 | CD4 | CD8 | CD163 | EpCAM | CD45 | CD3 | CD4 | CD8 | CD163 |
| 1 | 1.52 | 9.23 | 5.03 | 9.40 | 0.98 | 42.60 | 22.50 | 14.80 | 1.12 | 0.09 | 0.78 | 50.20 |

| | | | | | | | | | | | | |
|---|---|---|---|---|---|---|---|---|---|---|---|---|
| 2 | 0.28 | 0.16 | 0.03 | 0.06 | 0.06 | 0.46 | 6.31 | 2.50 | 0.40 | 0.03 | 0.60 | 74.10 |
| 3 | 4.08 | 2.74 | 2.10 | 5.94 | 2.82 | 3.40 | 15.40 | 14.80 | 1.30 | 0.90 | 2.84 | 83.80 |
| 4 | 1.12 | 4.75 | 2.17 | 5.31 | 0.96 | 5.54 | 1.65 | 6.24 | 0.21 | 0.00 | 1.25 | 73.10 |
| 5 | 9.16 | 10.50 | 11.00 | 30.90 | 9.83 | 20.90 | 4.88 | 47.40 | 1.37 | 1.30 | 0.45 | 88.80 |
| 6 | 0.00 | 7.53 | 13.50 | 20.40 | 1.54 | 8.70 | 57.30 | 20.40 | 0.34 | 0.19 | 0.35 | 91.90 |
| 7 | 0.71 | 22.90 | 8.53 | 7.49 | 2.21 | 34.90 | 1.86 | 47.20 | 1.25 | 0.69 | 0.83 | 94.50 |
| 8 | 0.34 | 1.12 | 1.04 | 1.01 | 0.58 | 1.71 | 3.12 | 14.70 | 0.30 | 0.09 | 0.49 | 93.30 |
| 9 | 1.43 | 3.52 | 12.50 | 19.70 | 6.49 | 5.28 | 4.99 | 49.00 | 12.50 | 2.76 | 0.50 | 94.90 |
| 10 | 6.94 | 50.60 | 2.96 | 7.12 | 6.91 | 69.10 | 0.80 | 60.70 | 0.21 | 2.63 | 5.85 | 89.10 |
| 11 | 10.40 | 15.30 | 6.13 | 9.64 | 4.63 | 19.90 | 12.70 | 72.30 | 1.29 | 1.73 | 0.80 | 98.50 |
| 12 | 0.62 | 3.96 | 0.74 | 6.85 | 2.18 | 13.90 | 9.76 | 1.22 | 0.07 | 0.03 | 0.07 | 69.80 |
| 13 | 1.51 | 1.88 | 1.02 | 0.77 | 1.69 | 10.80 | 13.60 | 29.70 | 4.75 | 6.92 | 20.90 | 92.80 |
| 14 | 1.00 | 6.71 | 6.59 | 6.58 | 2.11 | 23.70 | 2.64 | 8.27 | 0.16 | 0.10 | 0.16 | 59.50 |
| 15 | 0.00 | 8.75 | 12.00 | 18.60 | 9.97 | 4.95 | 1.80 | 14.50 | 0.23 | 0.11 | 0.86 | 79.80 |
| 16 | 0.00 | 7.59 | 9.15 | 16.10 | 11.60 | 2.52 | 10.00 | 50.30 | 0.89 | 0.86 | 0.33 | 89.00 |
| 17 | 0.00 | 2.60 | 4.11 | 8.86 | 3.79 | 5.43 | 0.00 | 15.30 | 0.58 | 0.28 | 8.00 | 60.50 |
| 18 | 2.46 | 8.16 | 15.20 | 2.33 | 18.70 | 3.42 | 4.94 | 14.40 | 15.60 | 14.20 | 6.90 | 90.00 |
| 19 | 2.05 | 4.53 | 20.60 | 1.92 | 14.00 | 4.34 | 5.03 | 11.30 | 6.03 | 33.70 | 10.20 | 91.70 |
| 20 | 6.31 | 55.60 | 9.91 | 1.94 | 11.30 | 7.19 | 1.48 | 57.60 | 0.94 | 7.28 | 13.80 | 95.50 |
| 21 | 2.50 | 0.65 | 6.92 | 1.77 | 4.28 | 64.90 | 1.73 | 2.47 | 0.68 | 64.60 | 4.08 | 2.80 |
| 22 | 9.32 | 3.22 | 30.30 | 10.40 | 72.10 | 11.00 | 14.00 | 3.99 | 0.39 | 1.60 | 2.50 | 39.20 |
| 23 | 0.00 | 3.94 | 11.60 | 15.00 | 3.85 | 2.62 | 9.09 | 86.70 | 1.56 | 31.80 | 1.09 | 87.50 |
| 24 | 0.00 | 0.20 | 0.28 | 1.91 | 0.10 | 0.86 | 0.00 | 27.20 | 0.86 | 6.01 | 1.42 | 75.90 |
| 25 | 0.00 | 0.31 | 3.04 | 4.03 | 3.04 | 1.43 | 3.85 | 3.14 | 0.33 | 25.90 | 1.81 | 77.10 |
| 26 | 0.97 | 2.28 | 1.71 | 2.13 | 1.66 | 2.63 | 8.04 | 78.80 | 2.33 | 47.50 | 0.37 | 94.40 |
| 27 | 0.16 | 0.45 | 0.79 | 1.03 | 1.56 | 3.70 | 2.20 | 25.30 | 2.12 | 12.80 | 3.12 | 77.80 |
| 28 | 0.00 | 0.27 | 1.84 | 0.62 | 0.21 | 0.26 | 5.62 | 27.60 | 1.84 | 17.10 | 0.84 | 75.30 |
| 29 | 0.22 | 2.88 | 1.28 | 0.43 | 0.00 | 4.63 | 3.64 | 19.00 | 7.97 | 18.30 | 3.12 | 94.20 |
| 30 | 3.93 | 6.70 | 3.36 | 14.60 | 1.92 | 7.98 | 5.62 | 33.70 | 0.85 | 1.96 | 1.04 | 90.60 |
| 31 | 1.37 | 2.42 | 18.60 | 5.54 | 10.50 | 4.14 | 3.99 | 18.90 | 1.85 | 8.35 | 4.10 | 84.20 |
| 32 | 9.09 | 2.22 | 35.50 | 43.60 | 27.60 | 2.77 | 81.80 | 12.70 | 0.94 | 7.31 | 2.09 | 76.90 |

| | | | | | | | | | | | | |
|---|---|---|---|---|---|---|---|---|---|---|---|---|
| 33 | 0.35 | 6.58 | 16.70 | 15.60 | 9.02 | 11.40 | 12.70 | 20.50 | 3.65 | 6.36 | 8.03 | 75.10 |
| 34 | 3.97 | 29.70 | 60.30 | 68.00 | 22.90 | 5.53 | 7.68 | 27.80 | 0.53 | 0.43 | 0.26 | 94.20 |
| 35 | 1.81 | 6.08 | 8.05 | 20.80 | 10.80 | 47.10 | 0.97 | 21.50 | 0.78 | 1.37 | 1.08 | 84.20 |
| 36 | 1.55 | 4.06 | 25.20 | 11.80 | 24.00 | 17.20 | 4.80 | 46.20 | 3.91 | 4.28 | 4.78 | 92.30 |
| 37 | 0.94 | 2.10 | 3.02 | 3.77 | 1.78 | 4.47 | 1.70 | 38.00 | 1.55 | 14.50 | 2.52 | 86.00 |
| 38 | 0.96 | 13.40 | 9.53 | 4.84 | 8.36 | 5.62 | 1.03 | 14.20 | 0.47 | 9.87 | 1.15 | 71.60 |
| 39 | 0.57 | 0.33 | 7.57 | 4.82 | 4.52 | 0.00 | 15.20 | 9.04 | 1.11 | 4.90 | 2.17 | 47.80 |
| 40 | 2.22 | 3.70 | 14.70 | 8.17 | 16.80 | 9.78 | 18.80 | 32.60 | 3.09 | 47.20 | 3.25 | 98.90 |
| 41 | 0.00 | 0.25 | 5.93 | 18.00 | 11.80 | 14.00 | 64.30 | 50.30 | 1.44 | 12.10 | 3.14 | 94.40 |
| 42 | 4.53 | 7.97 | 8.11 | 11.10 | 4.95 | 10.20 | 8.42 | 40.50 | 1.23 | 4.25 | 0.60 | 95.50 |
| 43 | 0.57 | 0.33 | 7.57 | 4.82 | 4.52 | 0.00 | 15.20 | 9.04 | 1.11 | 4.90 | 2.17 | 47.80 |
| 44 | 3.17 | 3.76 | 2.73 | 10.50 | 3.02 | 9.39 | 69.90 | 23.60 | 0.53 | 1.53 | 0.47 | 89.90 |
| 45 | 1.06 | 1.50 | 12.80 | 1.30 | 2.95 | 3.27 | 4.06 | 9.16 | 0.99 | 0.96 | 0.78 | 62.00 |
| 46 | 1.81 | 1.15 | 16.00 | 49.40 | 2.07 | 5.01 | 14.90 | 6.71 | 0.06 | 1.49 | 5.14 | 88.80 |
| 47 | 0.36 | 0.27 | 15.40 | 16.80 | 1.37 | 0.41 | 0.97 | 36.20 | 1.47 | 8.40 | 6.36 | 91.50 |
| 48 | 0.35 | 1.40 | 37.20 | 11.40 | 2.56 | 0.00 | 16.00 | 4.03 | 1.35 | 2.98 | 3.48 | 50.00 |
| 49 | 0.00 | 0.92 | 3.12 | 3.20 | 0.78 | 15.90 | 15.00 | 5.39 | 0.84 | 2.94 | 0.78 | 64.20 |
| 50 | 1.35 | 3.30 | 3.90 | 3.26 | 4.78 | 0.88 | 3.59 | 15.20 | 0.78 | 7.76 | 1.00 | 84.60 |
| 51 | 2.62 | 4.32 | 7.55 | 5.31 | 8.56 | 16.90 | 5.42 | 5.58 | 3.20 | 3.91 | 34.80 | 62.70 |
| 52 | 41.70 | 40.20 | 5.19 | 3.39 | 3.83 | 66.50 | 39.90 | 59.80 | 0.62 | 17.30 | 2.45 | 91.70 |
| 53 | 0.00 | 0.54 | 0.49 | 0.31 | 0.16 | 4.26 | 4.00 | 19.60 | 0.73 | 0.62 | 0.49 | 76.60 |
| 54 | 4.85 | 1.52 | 3.61 | 2.51 | 2.70 | 3.44 | 0.24 | 6.13 | 0.27 | 2.17 | 0.31 | 88.90 |

| **Specimen #** | **Galectin 9** | | | | | **CEACAM1** | | | | |
|---|---|---|---|---|---|---|---|---|---|---|
| | EpCAM | CD45 | CD3 | CD8 | CD163 | EpCAM | CD45 | CD3 | CD8 | CD163 |
| 1 | 68.90 | 43.80 | 22.50 | 20.10 | 82.90 | 65.30 | 10.60 | 10.40 | 7.49 | 17.90 |
| 2 | 29.40 | 15.70 | 0.38 | 0.42 | 86.80 | 68.90 | 3.13 | 1.84 | 0.23 | 25.80 |
| 3 | 95.20 | 72.10 | 25.70 | 24.30 | 95.60 | 65.60 | 14.00 | 14.90 | 17.40 | 50.10 |
| 4 | 51.10 | 41.40 | 10.50 | 5.06 | 86.90 | 0.31 | 4.72 | 2.22 | 0.72 | 10.30 |
| 5 | 91.30 | 92.00 | 19.70 | 11.20 | 97.80 | 56.90 | 7.56 | 4.94 | 3.72 | 11.20 |
| 6 | 58.30 | 34.40 | 61.10 | 60.40 | 83.90 | 66.00 | 2.87 | 0.23 | 0.45 | 4.69 |

| | | | | | | | | | | |
|---|---|---|---|---|---|---|---|---|---|---|
| 7 | 36.60 | 63.80 | 4.45 | 24.90 | 85.90 | 83.70 | 17.70 | 2.31 | 14.80 | 15.80 |
| 8 | 92.70 | 13.00 | 1.18 | 1.57 | 88.30 | 94.10 | 12.80 | 0.64 | 1.34 | 75.60 |
| 9 | 87.60 | 50.20 | 3.12 | 4.56 | 95.00 | 72.40 | 8.68 | 0.66 | 0.56 | 13.60 |
| 10 | 96.10 | 77.30 | 51.60 | 61.80 | 87.40 | 81.70 | 67.40 | 54.70 | 64.50 | 83.10 |
| 11 | 7.20 | 80.80 | 3.97 | 3.96 | 95.60 | 73.80 | 19.80 | 2.11 | 1.74 | 22.40 |
| 12 | 93.40 | 3.51 | 2.18 | 0.92 | 45.90 | 83.70 | 0.21 | 0.04 | 0.04 | 9.58 |
| 13 | 90.30 | 28.50 | 8.07 | 12.10 | 77.10 | 95.60 | 6.86 | 2.76 | 6.04 | 33.90 |
| 14 | 93.60 | 17.50 | 3.03 | 4.71 | 92.30 | 38.20 | 0.39 | 0.28 | 0.63 | 6.51 |
| 15 | 63.20 | 7.25 | 0.52 | 1.02 | 82.00 | 95.70 | 1.20 | 0.32 | 2.12 | 8.65 |
| 16 | 76.30 | 41.70 | 7.56 | 5.69 | 93.00 | 87.40 | 10.90 | 1.35 | 2.66 | 24.80 |
| 17 | 90.30 | 28.50 | 7.07 | 12.10 | 77.10 | 95.60 | 6.86 | 2.76 | 6.04 | 33.90 |
| 18 | 93.00 | 41.30 | 74.00 | 78.50 | 92.40 | 84.90 | 1.62 | 15.00 | 26.20 | 4.21 |
| 19 | 93.20 | 30.10 | 70.50 | 84.90 | 92.10 | 90.30 | 1.38 | 5.00 | 5.92 | 2.46 |
| 20 | 93.90 | 57.30 | 86.60 | 87.10 | 89.40 | 85.00 | 10.90 | 89.50 | 68.60 | 11.50 |
| 21 | 83.00 | 4.24 | 15.90 | 16.10 | 17.80 | 95.50 | 1.27 | 9.52 | 7.14 | 2.54 |
| 22 | 25.50 | 29.10 | 36.40 | 41.90 | 47.90 | 98.10 | 2.69 | 5.32 | 7.17 | 14.10 |
| 23 | 25.00 | 95.90 | 78.30 | 85.70 | 99.10 | 22.50 | 0.86 | 0.00 | 9.52 | 1.94 |
| 24 | 86.90 | 56.50 | 50.00 | N. A. | 86.10 | 0.00 | 0.08 | 50.00 | N.A. | 0.43 |
| 25 | 76.20 | 30.00 | 57.10 | 72.20 | 92.90 | 71.40 | 1.42 | 21.40 | 22.20 | 5.28 |
| 26 | 35.80 | 62.70 | 52.90 | 65.50 | 97.70 | 20.20 | 0.23 | 0.00 | 0.00 | 0.46 |
| 27 | 10.70 | 28.60 | 20.50 | 30.00 | 48.30 | 58.40 | 0.90 | 0.00 | 0.00 | 2.21 |
| 28 | 95.70 | 33.60 | 36.40 | 42.90 | 93.50 | 65.20 | 0.04 | 0.00 | 0.00 | 0.09 |
| 29 | 17.40 | 16.50 | 55.60 | 42.90 | 83.00 | 14.60 | 0.00 | 11.10 | 0.00 | 0.00 |
| 30 | 92.00 | 52.50 | 75.30 | 84.25 | 94.90 | 95.00 | 4.78 | 39.00 | 44.70 | 5.81 |
| 31 | 87.20 | 42.20 | 86.40 | 86.30 | 96.20 | 70.90 | 1.92 | 5.29 | 8.52 | 2.38 |
| 32 | 4.95 | 53.60 | 78.10 | 89.20 | 94.20 | 10.70 | 3.57 | 1.37 | 1.92 | 4.23 |
| 33 | 77.80 | 61.20 | 80.40 | 78.80 | 83.30 | 85.70 | 13.60 | 20.10 | 32.50 | 8.64 |
| 34 | 84.80 | 36.60 | 90.70 | 92.00 | 93.70 | 7.62 | 0.48 | 1.69 | 2.30 | 0.62 |
| 35 | 19.80 | 51.60 | 55.70 | 86.50 | 88.50 | 20.70 | 5.20 | 17.70 | 19.90 | 3.07 |
| 36 | 96.70 | 44.10 | 77.80 | 94.60 | 89.40 | 31.20 | 3.77 | 14.90 | 11.90 | 3.99 |
| 37 | 84.90 | 46.90 | 88.90 | 92.10 | 94.50 | 99.40 | 85.70 | 95.20 | 92.10 | 99.40 |

| | | | | | | | | | | |
|---|---|---|---|---|---|---|---|---|---|---|
| 38 | 89.30 | 33.90 | 88.60 | 80.00 | 73.30 | 73.10 | 1.11 | 4.55 | 9.41 | 1.72 |
| 39 | 40.40 | 97.30 | 95.00 | 96.30 | 98.50 | 87.90 | 5.51 | 11.80 | 15.50 | 3.27 |
| 40 | 41.60 | 39.30 | 71.20 | 72.60 | 93.70 | 85.90 | 1.82 | 12.90 | 17.70 | 3.20 |
| 41 | 75.00 | 90.20 | 98.70 | 99.40 | 97.50 | 22.50 | 0.43 | 0.00 | 0.00 | 0.49 |
| 42 | 76.60 | 29.10 | 56.00 | 66.10 | 61.40 | 30.00 | 1.99 | 8.97 | 7.21 | 1.07 |
| 43 | 71.10 | 2.39 | 10.30 | 17.60 | 19.40 | 75.40 | 1.38 | 8.04 | 14.70 | 7.29 |
| 44 | 90.30 | 66.10 | 82.80 | 85.00 | 85.60 | 14.00 | 0.96 | 2.72 | 2.77 | 2.39 |
| 45 | 93.30 | 48.30 | 88.10 | 90.40 | 96.70 | 34.10 | 1.22 | 3.68 | 5.16 | 3.41 |
| 46 | 78.40 | 15.50 | 52.90 | 65.10 | 75.70 | 38.40 | 1.44 | 3.23 | 2.33 | 10.70 |
| 47 | 81.30 | 60.40 | 37.90 | 68.90 | 88.60 | 88.20 | 4.35 | 1.72 | 2.22 | 1.90 |
| 48 | 66.20 | 11.00 | 35.70 | 45.60 | 84.20 | 41.30 | 4.71 | 5.71 | 2.94 | 4.85 |
| 49 | 96.10 | 34.10 | 59.10 | 92.00 | 93.90 | 28.10 | 1.92 | 8.89 | 9.09 | 5.76 |
| 50 | 45.80 | 92.70 | 96.50 | 94.70 | 99.40 | 96.80 | 10.60 | 38.30 | 72.70 | 40.00 |
| 51 | 97.80 | 78.10 | 96.00 | 100.00 | 95.00 | 91.60 | 11.90 | 73.30 | 85.70 | 21.20 |
| 52 | 51.90 | 84.90 | 87.80 | 95.40 | 90.40 | 89.10 | 17.20 | 88.60 | 58.30 | 23.40 |
| 53 | 67.40 | 27.70 | 84.70 | 53.10 | 98.90 | 77.60 | 2.29 | 10.50 | 13.50 | 2.00 |
| 54 | 40.70 | 18.40 | 81.00 | 71.70 | 95.40 | 59.20 | 1.22 | 46.90 | 45.80 | 2.25 |
