## Supplemental Table 2 for "TIM3^+^ Tumor Associated M2 Macrophages Impair Antitumor T Cell Immunity and Promote Gastric Cancer Progression and Peritoneal Metastasis"

**Supplemental Table 2- KEY RESOURCES**
**Reagents, Kits and other resources used in this study**

| Reagents | Source | Identifier |
|---|---|---|
| LIVE/Dead Fixable Aqua Dead Cell Stain Kit | Life Techgologiest | L34965 |
| BD Cytofix/Cytoperm Plus | BD Biosciences | 555028 |
| Foxp3/Transcription Factor Staining Buffer Set | eBiosciences | 5523-00 |
| rhIL-2 | R&D Systems | BT-002-10 |
| rhCCL20 (MIP3a) | R&D Systems | 360-MP-025/CF |
| Mouse M-CSF Recombinant Protein | PeproTech | 315-02-2UG |
| Mitomycin C | Thermo Scientific | J63193 |
| Ravoxertinib (GDC-0994) | Selleck Chemicals | S7554 |
| BI-D1870 | Selleck Chemicals | S2843 |
| Matrigel Matrix | Corning | 356237 |
| Standard Macrophage Depletion Kit | Encapsula Nano Sciences | CLD-8901 |
| Dispase | Gibco | 17105-041 |
| Collagenase IV | Gibco | 9001-12-1 |
| Luna Script SuerMix | New England BioLabs | M3010L |
| Applied Biosystems | PwerUp SYBR Green Master Mix | 100029284 |
| iBlot 3 Transfer Stacks, midi, PVDF | invitrogen | IB34001 |
| SurePAGE, Bis-Tris, 10x8 4-12%, 12 wells | Genscript USA | M000653 |
| Mouse CCL20/MIP-3 alpha Quantikine ELISA Kit | R&D Systems | DY360-05 |
| Human Cytokine Antibody Array C5 | RayBiotech | AAH-5-8 |
| Proteome Profiler Human Phospho-Kinase Array Kit | R&D | ARY003C |

### Quantitative RT-PCR primers used in this study

| Primer name | Oligo nucleotide sequence |
|---|---|
| CCL1-F | CCCCTTCAGCGACTAGAGAG |
| CCL1-R | TCTTGGGGTCGGCACAGAT |
| CCL2-F | TCTGTGCCTGCTGCTCATAG |
| CCL2-R | AGATCTCCTTGGCCACAATG |
| CCL4-F | TGCTAGTAGCTGCCTTCTGC |
| CCL4-R | GCTTCCTCGCGGTGTAAGAA |
| CCL5-F | CCAGCAGTCGTCTTTGTCAC |
| CCL5-R | CTCTGGGTTGGCACACACTT |
| CCL7-F | ATCCCTAAGCAGAGGCTGGA |
| CCL7-R | GTCCTGGACCCACTTCTGTG |
| IL-1beta-F | CCACCTCCAGGGACAGGATA |
| IL-1beta-R | AACACGCAGGACAGGTACAG |
| IL-6-F | CGGACAGCTTGAACAGAATGT |
| IL-6-R | ACCATCCCACTCACACCTCA |
| MIP-3alpha(CCL20)-F | GCGAATCAGAAGCAGCAAGC |
| MIP-3alpha(CCL20)-R | TTGGATTTGCGCACACAGAC |
| IL-10-F | TCAAGGCGCATGTGAACTCC |
| IL-10-R | GATGTCAAACTCACTCATGGCT |
| GAPDH-5 | ACCCAGAAGACTGTGGATGG |

| | |
|---|---|
| GAPDH-3 | TCTAGACGGCAGGTCAGGTC 3 |
| Tim3-F | TTGGACATCCAGATACTGGCT |
| Tim3-R | GGTGGTAAGCATCCTTGGAA |
| Galectin-9F | TTCTCCCAGCCTGTCTGTTT |
| Galectin-9R | TGGCGGGAGTAGAGAACATC |
| CEACAM1-F | CAGTGACCCAGTCACCTTGA |
| CEACAM1-R | GCCAGGAGTACTGTGCAGGT |
| mArginase1-F | CTCCAAGCCAAAGTCCTTAGAG |
| mArginase1-R | AGGAGCTGTCATTAGGGACATC |
| mGAPDH-F | CATCACTGCCACCCAGAAGACTG |
| mGAPDH-R | ATGCCAGTGAGCTTCCCGTTCAG |
| mCCL2-F | AAGGCAAAGGCAGTTCTCAA |
| mCCL2-R | GGGAGGGCAGGTAGAAAAAG |
| mIL-10-F | CCAAGCCTTATCGGAAATGA |
| mIL-10-R | TTTTCACAGGGGAGAAATCG |
| mTIM3-F | TTGGAGTGGGAGTCTCTGCT |
| mTIM3-R | GGCAAGTTGGCCAGTGTAAT |
| mGalectin9-F | ATTCCAAATGGGCTTTACCC |
| mGalectin9-R | AGGTGGAAAGCAATGTCACC |

**Antibodies used for Flow cytometry, CyTOF, WB and co-IF**

| Antibodies | Source | Identifier |
|---|---|---|
| FITC-anti-human CD45 | BioLegend | 304006 |
| APC-Cy7 anti-human CD3 | BD Biosciences | 557832 |
| Alexa Fluor700 anti-human CD8 | BD Biosciences | 557945 |
| PerCP-Cy5.5 anti-human CD4 | BioLegend | 357414 |
| PE anti human-CD366 (Tim-3) | BioLegend | 345005 |
| APC-anti-human EpCAM | BioLegend | 324207 |
| BV421 anti-human CD66a/c/e (CEACAM1) | BioLegend | 342313 |
| PerCP-Cy5.5 anti human Galectin-9 | BioLegend | 348909 |
| PE-CF594 anti-human CD279 (PD1) | BD Biosciences | 566846 |
| PE-Cy7 anti-human CD274 (PD-L1) | BioLegend | 374505 |
| APC anti human-TNF-$\alpha$ | BioLegend | 502913 |
| PE-Cy7 anti-human IFN-$\gamma$ | BioLegend | 506518 |
| PE-Cy5 anti-humanCD163 | BioLegend | 333644 |
| PerCP-Cy5.5 anti-human CD163 | BD Biosciences | 563887 |
| FITC anti-human CD206 | BioLegend | 321103 |
| BV421 anti-human Granzyme B | BD Biosciences | 563389 |
| PE-CF594 anti-human Perforin | BD Biosciences | 5093012 |
| anti-human CD279 (PD-1) | BioXCell, InVivoMab | BE0188 |
| anti-human CD366 (Tim-3) | BioLegend | 345038 |
| Purified mouse IgG1 $\kappa$ | BioLegend | 401402 |
| anti-h/m/rb-Actin | R&D | 937216 |

| | | |
|---|---|---|
| Anti-mouse IgG HRP-linked Antibody | Cell Signaling | 7076 |
| Anti-rabbit IgG HRP-linked Antibody | Cell Signaling | 7074 |
| TIM-3/HAVCR2 antibody | SinoBiological | 51152-T48 |
| anti-human CD196 (CCR6) | eBioSciences, invitrogen | 14-1969-80 |
| Akt antibody | Cell Signaling | 9272 |
| Phospho-Akt Rabbit mAb | Cell Signaling | 4060 |
| Phospho-p44/42 MAPK (Erk1/2) | Cell Signaling | 4370 |
| RSK1/RSK2/RSK3 Rabbit mAB | Cell Signaling | 9355 |
| Phospho-p90RSK (Ser380) Rabbit mAb | Cell Signaling | 11989 |
| TIM-3 XP Rabbit mAb | Cell Signaling | 45208 |
| CD163 | Leica Biosystems | NCL-L-CD163 |
| YAP1 (63.7) | Santa cruz | sc-101196 |
| CD8 | Cell Signaling | 98941 |
| Ki-67 Ab-4 Rabbit | Fisher Scientific | Epredia RM9106S0 |
| APC-Cy7 anti-mouse CD3 | BioLegend | 100222 |
| Alexa Fluor700 anti-mouse CD8 | BioLegend | 126618 |
| PE anti-mouse IFNγ | BioLegend | 505826 |
| PE-CF594 anti-mouse Perforin | BioLegend | |
| PE anti-mouse CD68 | BioLegend | 137013 |
| BV421 anti-mouse CD163 | BioLegend | 155309 |
| APC anti-mouse CD206 | BioLegend | 141708 |
| PerCP anti-mouse F4/80 | BioLegend | 123125 |
| anti-mouse TIM-3 (CD366) | BioXCell, InVivoMab | BE01115 |
| anti-mouse PD-1 (CD279) | BioXCell, InVivoMab | BE0146 |
| anti-mouse PD-L1 | BioXCell, InVivoMab | BE0101 |
